# Structural insights into the molecular recognition of integrin αVβ3 by RGD-containing ligands: The role of the specificity-determining loop (SDL)

**DOI:** 10.1101/2024.09.23.614545

**Authors:** Charles Mariasoosai, Santanu Bose, Senthil Natesan

## Abstract

Integrin αVβ3 is a prominent member of the “RGD-recognizing” integrin family of cell surface receptors. αVβ3 binds to various extracellular matrix (ECM) proteins and oxysterols such as 25-hydroxycholesterol, is implicated in several diseases, including cancer metastasis, lung fibrosis, inflammation, and autoimmune diseases, and is pursued as a valuable therapeutic target. Despite enormous efforts to seek a pure antagonist, to date, no single drug candidate has successfully reached clinics due to associated partial agonism and toxicity issues. Developing effective and safe inhibitors require a thorough understanding of the molecular interactions and structural changes related to the receptor’s activation and inhibition mechanisms. This study offers a comprehensive residue-residue contact and network analyses of the ligand-binding β-propeller βI domains (headpiece) based on all available experimental structures of integrin αVβ3 in unliganded, agonist-, antagonist-, and antibody-bound states. The analyses reveal many critical interactions that were not reported before and show that specific orientation and interactions of residues from the specificity-determining loop (SDL) are critical in molecular recognition and regulation. Also, the network analysis reveals that residues from the nearby allosteric site (site II) connect to the primary RGD-binding site via SDL, which likely acts as an interface between the two sites. Our results provide valuable insights into molecular interactions, structural changes, distinct features of the active and inactive headpiece conformations, the role of SDL in ligand recognition, and SDL-mediated allostery. Thus, the insights from this study may facilitate the designing of pure antagonists or site II-mediated allosteric modulators to integrin αVβ3 to treat various diseases.

## INTRODUCTION

Integrins are bidirectional signaling molecules mediating cell-matrix and cell-cell adhesion and participating in various intracellular signaling pathways[1]. Integrins are heterodimers assembled from eighteen α and eight β subunits (resulting in 24 unique heterodimer combinations in humans), allowing them to interact with a wide variety of extracellular matrix (ECM) ligands[2]. The ability of the cell to adapt to the changes in both extracellular and intracellular environments is mediated by multi-directional integrin signaling (inside-out, outside-in, and inside-in)[3–5]. The association of integrins with several major diseases with unmet medical needs makes them one of the most attractive therapeutic targets. Currently, six integrin inhibitor drugs are in clinical use, and there have been at least ∼216 clinical trials related to integrin-based therapies (The clinicaltrials.gov. database was searched with the term ‘integrin’ on June 10, 2023). Among the integrin heterodimers, αVβ3 is one of the most studied members due to its role in inflammation, thrombosis, arthritis, glioblastoma, and cancer [6, 7] [8, 9]. Additionally, many viruses have been shown to utilize αVβ3 as a receptor or co-receptor for their cellular entry[10]. Furthermore, we recently demonstrated the direct participation of integrin αVβ3 in triggering a proinflammatory response through oxysterols during virus infections.

Integrin αVβ3 consists of αV and β3 subunits, and each subunit contains a larger ectodomain, single transmembrane helix, and a short cytoplasmic tail. In the ectodomain, the β-propeller domain of αV and the βI domain of β3 subunits associate non-covalently to form the headpiece that provides the primary binding interfaces for many ECM and other ligands. In the αV subunit, β-propeller is connected to the transmembrane helix via thigh, calf-1, and calf-2 domains. Similarly, in the β3 subunit, the βI domain is connected to the transmembrane helix via PSI, E1-4, and βT domains (**Fig. 1A**). Integrin αVβ3 has two ligand binding sites in the headpiece: the primary RGD-binding site (site I) and an allosteric site, herein referred to as site II. Numerous RGD-motif containing ECM proteins, such as fibrinogen, vitronectin, von Willebrand Factor, thrombospondin-1, fibrillin, tenascin, platelet endothelial cell adhesion molecule 1 (PECAM-1), and many other ligands bind to site I of integrin αVβ3[11] [12]. Similarly, chemokine fractalkine, secreted phospholipase A2 type IIA, CX3CL1, CXCL12, CD40L, and a few hydroxylated forms of cholesterol such as 25-hydroxycholesterol and 24-(S)-hydroxycholesterol have been shown to bind at site II and likely regulate the binding of ECM ligands at site I of integrin αVβ3 allosterically[13–17]. Site II is located adjacent but distal to the primary binding site at the interface between the β-propeller and the βI domains (**Fig. 1B**). Intriguingly, the two binding sites are connected by the specificity-determining loop (SDL) located near the sites. Many of the primary RGD and allosteric site ligands such as chemokine fractalkine, secreted phospholipase A2 type IIA, CX3CL1, CXCL12, CD40L, and 25HC were reported to interact with the specificity determining loop[13–16, 18].

**Figure 1.**
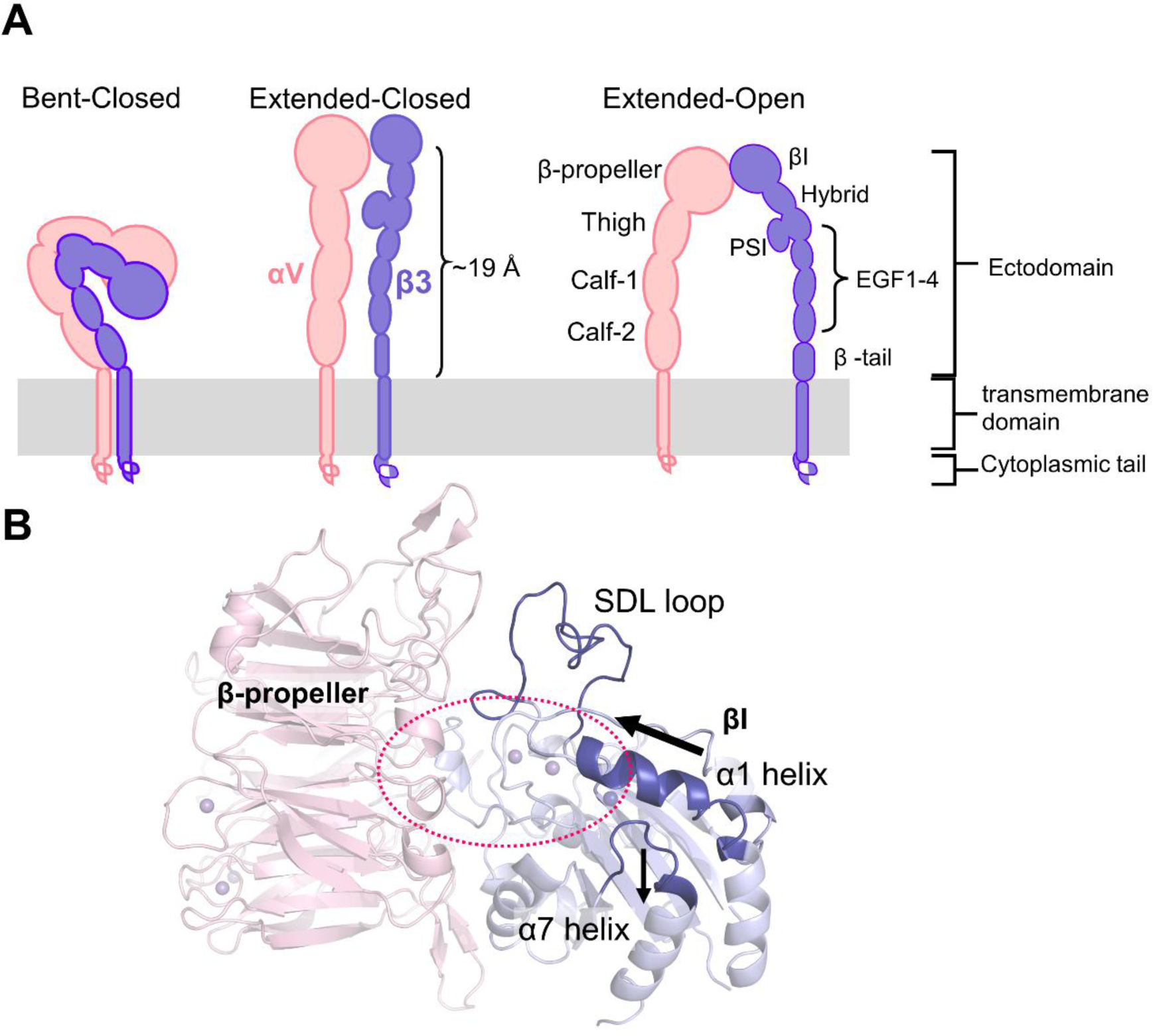
Structure and conformational states of αVβ3 integrin. **A)** The schematic diagram of the different conformational states. Each subunit of the αVβ3 heterodimer has a large ectodomain, a single transmembrane helix, and a short cytoplasmic tail. The ectodomain of each subunit consists of distinct subdomains and, together, exist in bent-closed, extended-closed, and extended-open conformational states. In the extended-open conformation, the integrin ectodomain can elongate up to ∼19 Å to interact with extracellular ligands. **B)** The primary RGD-binding site is located at the interface between the β-propeller and βI domains, collectively termed the headpiece. The secondary structure representation of the headpiece in the “unliganded” state (PDB ID: 1JV2) shows the RGD binding site in a red dotted ellipse and secondary structure elements such as α1 helix and b6-α7 loop that undergo conformational changes upon the receptor activation. Upon receptor activation by ligands, the α1-helix moves toward the site, and the F-α7 loop moves away from the α1 helix (arrows indicate the direction of movements). The residues from the specificity-determining loop (SDL) have been shown to participate in ligand binding and are critical for specific functional outcomes.

**Figure 2.**
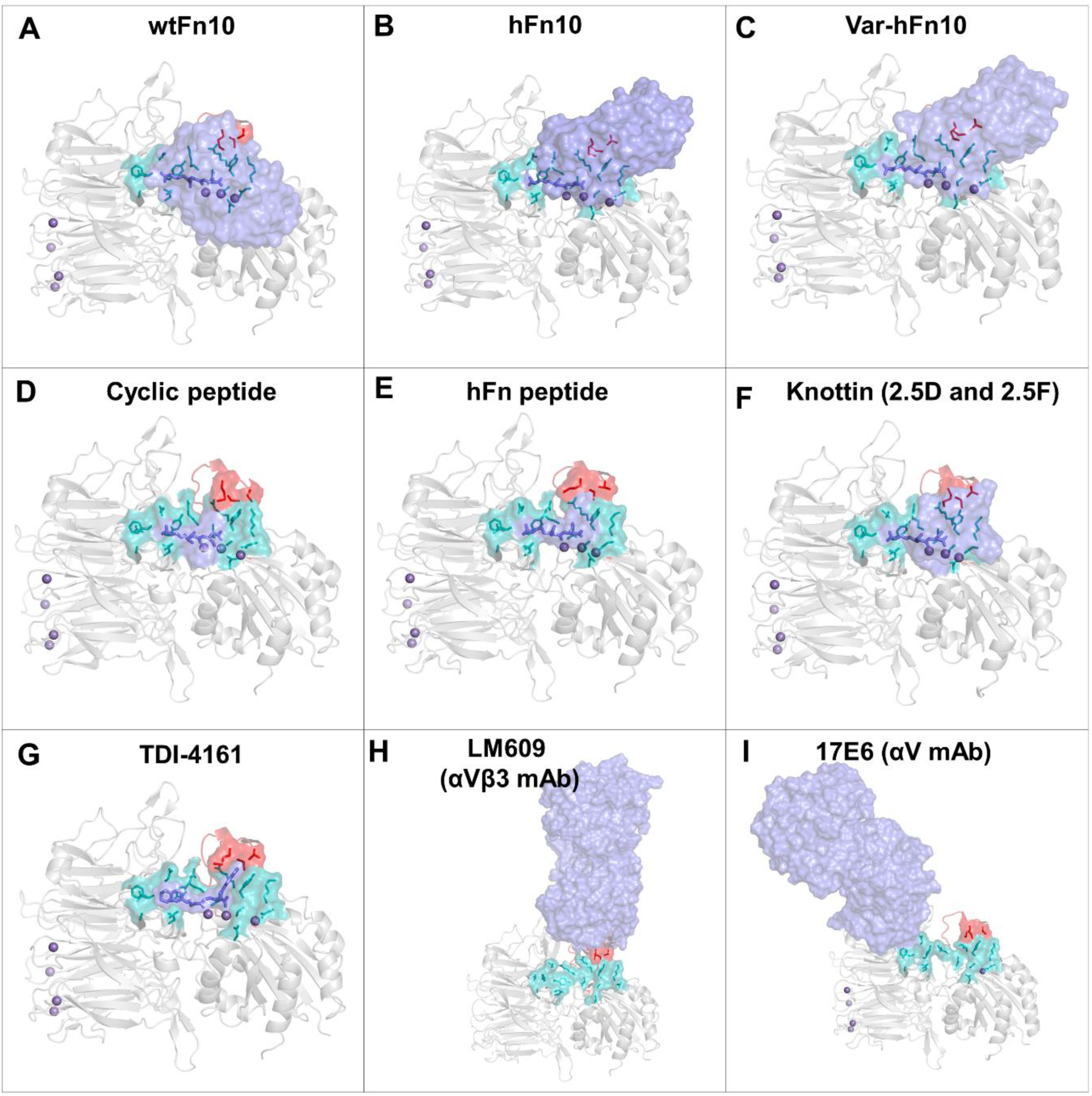
Experimental structures of the integrin αVβ3 headpiece bound to various ligands. The β-propeller and βI domains of the αVβ3 headpiece are shown in the cartoon secondary structure representation (gray), and the binding partners are shown in the surface representation (slate blue). In all the ligand-bound structures, the RGD motif binds in a similar orientation: The Asp (D) residue is oriented towards Mn^2+^ at the MIDAS site, and Arg (R) is oriented towards the β-propeller domain. The RGD motif is represented as sticks in dark blue. The primary binding site residues are shown in both stick (teal) and surface (cyan) representations. Residues from the specificity-determining loop (SDL) that interact with the ligands are shown in stick and surface representations in red. Mn^2+^ ions in the RGD binding site and the β-propeller domains were shown as purple spheres. **A)** wtFn10 is oriented towards the α7 helix of the βI domain. **B)** hFn10 binds in an entirely different orientation than wtFn10 and forms additional interactions with SDL of the βI domain. **C)** var-hFn10 binds in a similar orientation as hFn10. **D)** The cyclic RGD peptide. **E)** hFn10 peptide. **F)** Two knottin peptides, 2.5D and 2.5F, bind similarly despite their integrin subtype selectivity. **G)** TDI-4161, a small molecule inhibitor, binds like other RGD motif-containing ligands and mimics their orientation and interactions like hFn10. **H)** αVβ3-specific monoclonal antibody binds to the β-propeller residues and SDL loop of the βI domain, and **I)** αV-specific monoclonal antibody binds at the β-propeller domain. Both αV-specific and αVβ3-specific monoclonal antibodies bind at distinct sites and do not have any direct contact with the primary RGD binding site residues.

The outside-in activation of integrins refers to the shift from the bent-closed conformation to the extended-open conformation with a significantly increased affinity for ECM ligands [19] [20, 21]. In all the conformational states associated with the regulation of integrin, the non-covalent interactions within and between the βI and β-propeller domains of the headpiece seem intact and undergo subtle changes [22]. Thus, elucidating the conformational and structural changes within the integrin headpiece is critical to understand the integrin activation process[23–25]. The binding of ligands to integrin αVβ3 is a cation-dependent process during which two additional cations are acquired in and around the RGD binding site[26]. The ligand binding also induces specific structural changes as part of the receptor activation. The specific activation signatures include the movement of the α1 helix toward the RGD ligand binding site and the opening of the α7 loop[26, 27]. Several small-molecule integrin αVβ3 antagonists failed due to the formation of the ligand-induced binding site (LIBS) that resulted in the partial agonism of the receptor, likely causing fatal side effects[28, 29]. Currently, there is no single FDA-approved small molecule drug addressing the partial agonism of the receptor. The differences between the active and inactive conformations appear to be very subtle and extremely challenging to be accurately characterized while designing inhibitors. The mechanistic details of how the binding of ligands and their molecular interactions at the integrin headpiece, including site I and site II, and the role of the specificity-determining loop in affecting the specificity remain poorly understood. Thus, the lack of knowledge about the structural properties of the pure antagonists and associated structural and conformational changes warrants a thorough investigation of the agonist- and inhibitor-bound conformations of αVβ3 integrin. The availability of rich structural data with various agonists, antagonists, and antibodies makes integrin αVβ3 an ideal candidate for examining the critical molecular interactions in its various functional states[30].

In this study, we comprehensively analyzed the molecular interactions in the integrin αVβ3 headpiece in both active and inactive conformations. Specifically, we examined the differences in the secondary structural conformations and residue interactions at the RGD-binding site, site II, SDL, and metal ion coordination sites. Our results offer valuable insights into the interplay of the SDL conformational changes dictating ligand-specific molecular interactions with agonists and antagonists and their role in connecting the primary and allosteric binding sites. Learning from the agonist and antagonist-bound structures would enable us to understand the key differences in the molecular interactions and incorporate the inhibitory features in the design of novel small molecule therapeutics targeting integrin αVβ3.

## RESULTS AND DISCUSSION

### Overview of the experimental structures of integrin αVβ3

To unravel specific molecular interactions and conformational changes associated with the activation and inhibition of integrin αVβ3, we analyzed all the eighteen experimental structures available in the Protein Data Bank[31] (**Table 1**). Among the analyzed structures, six were in the *apo* “unliganded” form, and the remaining twelve were in the “liganded” form, bound to various agonists, antagonists, and antibodies. The liganded structures include a variety of ‘RGD’ motif-containing molecules, such as the wildtype tenth domain of fibronectin (wtFn10), its high-affinity mutant (hFn10), and another variant (var-hFn10), a cyclic RGD peptide (Cilengitide), and knottin peptides EETI 2.5D and 2.5F. Irrespective of the functional nature, i.e., either agonists (full or partial) or antagonists, all of the RGD ligands primarily bind to the receptor through their conserved RGD motif. The receptor was bound to a small-molecular inhibitor (TDI-5161) or two monoclonal antibodies (LM609 and 17E6) in the remaining structures.

**Table 1.**
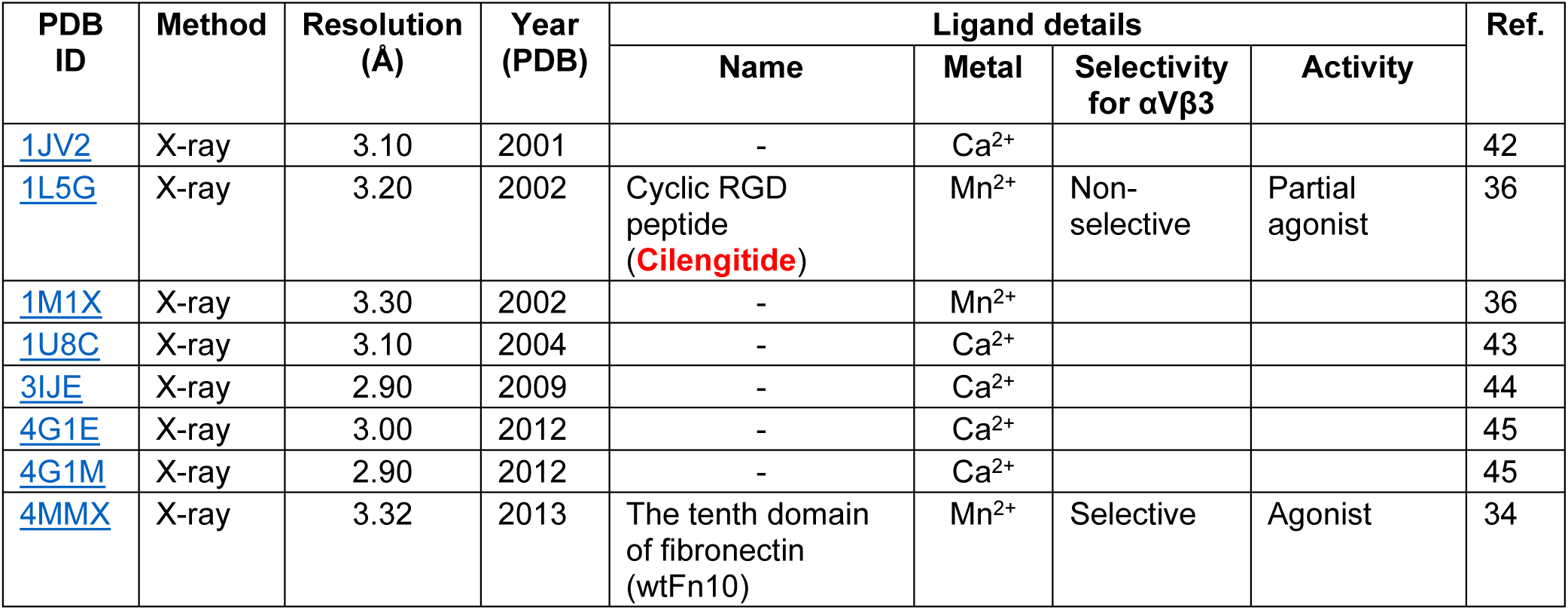
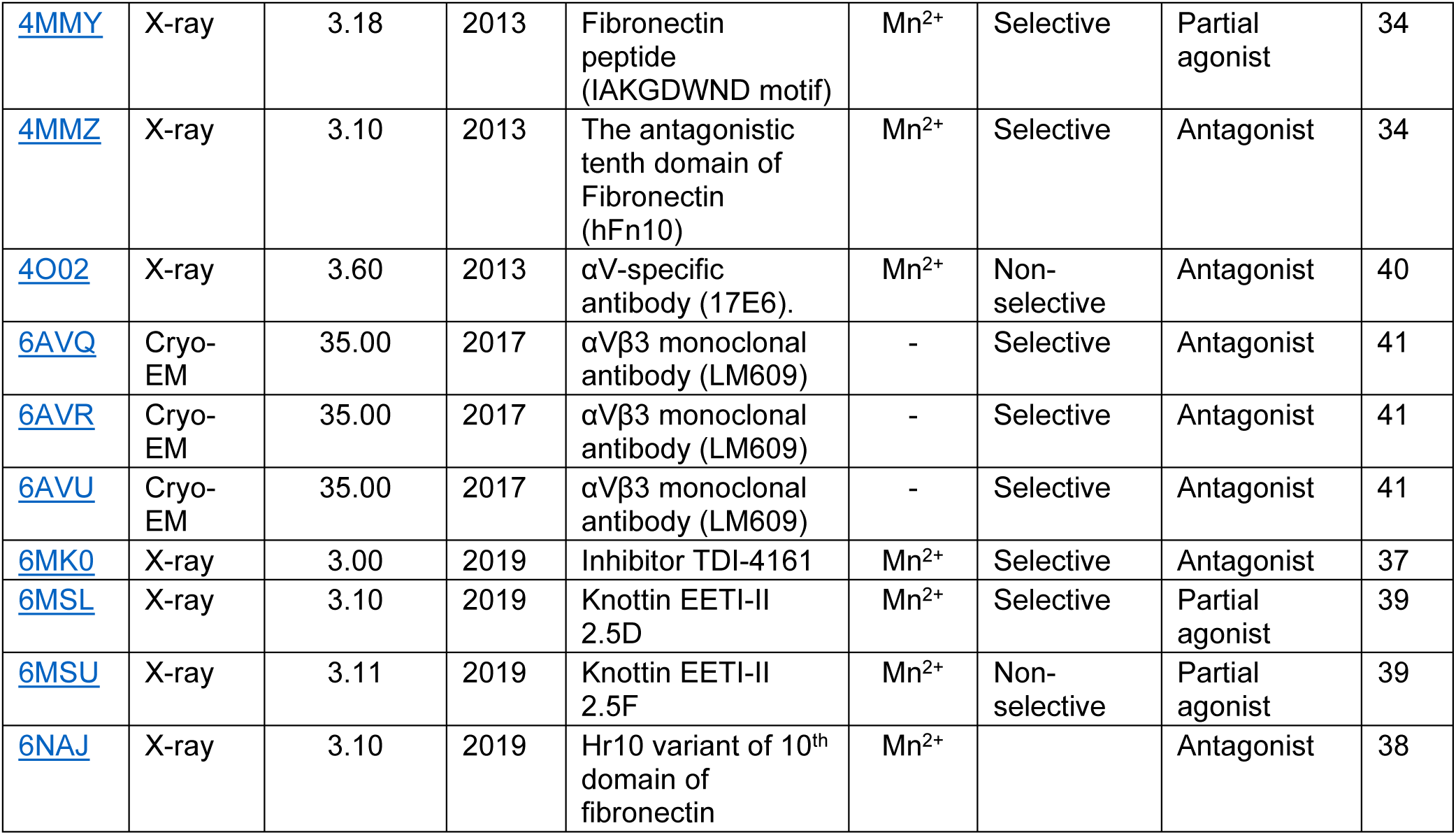
Experimental structures of integrin αVβ3 available in the Protein Data Bank.

The first “liganded” integrin αVβ3 structure was crystallized with a cyclic pentapeptide (Cilengitide, *RGDFV*, PDB ID:1L5G) that competitively inhibits various RGD-binding integrins, including integrins αVβ3 and α5β1. This crystal structure delineated the specific RGD-binding region in integrin αVβ3[26]. Later, the wtFn10-bound integrin αVβ3 complex (PDB ID: 4MMX) structure revealed the binding orientation and the associated conformational changes (activation signatures) induced by a natural ligand[27]. Surprisingly, a Ser1496Trp mutation in wtFn10 resulted in the high-affinity form of fibronectin (hFn10) that acts as a complete antagonist of the receptor (PDB ID: 4MMZ)[27]. In hFn10, the segment ^1492^PRGDWNEG^1499^ replaces ^1492^GRGDSPAS^1499^ of wtFn10. The mutations around the RGD segment of hFn10 from “**TP**RGDWN**E”** to “**IA**RGDWN**D”** caused the reorientation of Trp1496, which resulted in a higher affinity as compared to wtFn10, and produced partial agonism of the receptor (PDB ID: 4MMY)[27]. The higher affinity of hFn10 attained through the Ser1496Trp mutation and the orientation of Trp1496 that forms a π-π interaction with Tyr122 of the βI domain of the β3 subunit arrests the induction of the activation signal and retains the headpiece in an inactive state (liganded-inactive state)^34^. Attempts to facilitate a similar π-π interaction with Tyr122 by modifying the RGD-based small molecule antagonist of αVβ3, MK-429, resulted in the development of two new small molecule inhibitors, namely TDI-4161 and TDI-3761. These molecules exhibited antagonism similar to hFn10. As intended, TDI-4161 forms a similar π-π interaction with αVβ3[29].

Furthermore, studies were performed to gain antagonism against αIIbβ3, another member of the RGD-binding family of integrins. Superimposition of the crystal structures of αVβ3-hFn10 and αIIbβ3-eptifibatide (PDB ID: 2VDN) complexes shows that hFn10’s arginine of RGD cannot form any interaction with αIIbβ3’s Asp224, while the Ser1500 of hFn10 clashes with the αIIb’s β-propeller residues. Hence, the Ser1500 of hFn10 was mutated to glycine to avoid steric clashes, and arginine of RGD was substituted by homoarginine (HRG) with the extended alkyl chain by an additional methylene group, which facilitates the HRG sidechain to form a bidentate salt bridge with Asp224 of αIIb. As a result, the variant form of hFn10, var-hFn10, with “R” to “HRG” modification and S1500G mutation can inhibit αIIbβ3 as a pure antagonist. Though var-hFn10 has a lesser affinity towards αVβ3 than hFn10, the crystal structure of αVβ3-var-hFn10 complex (PDB ID: 6NAJ) shows that the var-hFn10 also induces similar conformational changes in αVβ3 headpiece to retain it in the “liganded-inactive*”* state[42]. In the case of engineered knottin mini proteins, EETI 2.5D and 2.5F were able to bind to αVβ3 and α5β1 with nanomolar affinities. The knottin mini proteins were primarily engineered by incorporating RGD motifs in the cysteine knot peptide from the squash family of protease inhibitors from *Ecballium elaterium* trypsin inhibitor (EETI). Even though these two knottins differ only by four residues, i.e., two residues preceding and succeeding the RGD motif (2.5D: GCP**QG<u>RGD</u>WA**PTSCKQDCLAGCVCGPNGGCG, 2.5F: GCP**RP<u>RGD</u>NP**PTSCKQDCLAGCVCGPNGGCG), knottin 2.5D binds selectively to αVβ3 while 2.5F binds to both αVβ3 and α5β1[43]. Crystal structures of knottins 2.5D (PDB ID: 6MSL) and 2.5F (PDB ID: 6MSU) bound to integrin αVβ3 show that both ligands bind in a similar orientation, and the difference in their selectivity was likely attributed to their flexibility at the binding site[44]. In addition, αVβ3 integrin was crystallized with two monoclonal antibodies: 17E6, which recognizes all αV-containing integrins, and LM-609, which is αVβ3-specific. The crystal structure of the 17E6 Fab–αVβ3 complex (PDB ID: 4O06) revealed that 17E6 binds to the β-propeller domain of the αV subunit, and its recognition site is distinct from the primary RGD-binding site[37]. In contrast, LM-609 binds at the interface between the β-propeller and βI domains near the RGD-binding site. Specifically, LM-609 interacts with the specificity-determining loop (SDL) but lacks direct contact with the primary RGD binding site residues. Furthermore, the LM-609 Fab-αVβ3 complex structure reveals that it inhibits the receptor through steric hindrance and binds to different conformations of αVβ3 such as closed (PDB ID: 6AVQ), open (PDB ID: 6AVR), and extended open conformations (PDB ID: 6AVU)[38]. Despite not being in direct contact with the primary RGD binding site, both 17E6 and LM-609 appear to block the site by steric hindrance.

Although wtFn10 and hFn10 differ only by a single mutation, Ser1496Trp, their binding to integrin αVβ3 results in opposite functional consequences. While wtFn10 induces conformational changes in the receptor towards its active state, hFn10-produced conformational changes arrest the receptor in an inactive state. Therefore, in most of the following analyses, we examined the differences in the molecular interactions and conformational changes between the wtFn10–αVβ3 and hFn10–αVβ3 complexes as prototypes for the liganded “active” and “inactive” states of integrin αVβ3.

### Major structural rearrangements associated with the binding of agonists and antagonists

Overall, careful analyses of the “unliganded” (PDB ID: 1JV2), “liganded-active” (PDB ID: 4MMX, wtFn10-αVβ3), and “liganded-inactive” conformations (PDB ID:4MMZ, hFn10 - αVβ3) of the integrin headpiece revealed that major ligand-induced conformational transformations occur mainly in the βI domain and only subtle changes in the β-propeller domain. Further, the variations in the β-propeller domain appear only at the residue sidechain interactions, whereas the backbone conformations remained unaffected in both liganded-active and inactive conformations^36^. Upon activation through the binding of agonists (for example, wtFn10), the βI domain undergoes two major structural rearrangements around the primary RGD-binding site: 1) the inward movement of α1 helix towards the RGD binding site and 2) the opening of the F-α7 loop and piston-like downward movement of α7-helix (**Fig. 1B**). In addition, subtle sidechain conformational changes occur in the SDL loop. Expectedly, the activation signatures observed in the “liganded-active” state were absent in the “liganded-inactive” structure. For example, the binding of antagonist hFn10 did not induce any conformational changes in and around the primary RGD-binding site that are part of the activation process. Remarkably, the activation signatures were arrested by hFn10. The conformation of α1 helix and F-a7 loop remain similar to the “unliganded” apo structure. Overall, analyses of the agonist (active) and antagonist (inactive) bound headpiece conformations show that αVβ3 antagonists such as hFn10 and small molecule inhibitor, TDI-4161, do not induce any conformational changes in the RGD-binding site and thereby retain the headpiece as the unliganded state [27, 29]. Interestingly, the SDL conformations show subtle differences in these structures. The secondary structure and overall conformation of SDL appear to be maintained by interactions between the residues within the loop and with residues present in the adjacent regions. These interactions appear to be altered distinctively by agonists and antagonists, suggesting that the changes in SDL are a critical part of the ligand recognition and binding processes.

### RGD motif interactions in the liganded-active and inactive state

The interactions of αVβ3 residues that primarily interact with the RGD ligands were analyzed to assess the similarity and differences among the agonists and antagonists (**Fig 3 and 4**). The RGD binding site is formed mainly by residues from the βI domain and extends to the β-propeller interface in the ectodomain headpiece (**Fig. 3A**). Notably, among the RGD motif residues, Arg (R) interacts with the β-propeller and Asp (D) interacts with the βI domain. Gly (G) is positioned in between and forms a nonpolar hydrophobic environment at the center of the RGD binding site. Both the β-propeller residues (Asp150, Phe177, Tyr178, Thr212, Ala213, Ala215, and Asp218) and βI domain residues (Tyr122, Ser123, Lys125, Asp126, Cys177, Met180, Thr182, Arg214, Asn215, Arg216, Asp251, Lys253) along with Mn^2+^ ion form the binding site and interact with the ligands (**Fig. 3B and 3C**). Among these residues, Asp150, Tyr178, Ala215, and Asp218 of the β-propeller domain and Tyr122, Ser123, Asn215, Arg216 of the βI domain, and Mn^2+^ at MIDAS (Metal ion-dependent adhesion site) interact with the RGD motif, irrespective of the ligand type and activity. In contrast to the macromolecule ligands, the small molecule inhibitor TDI-4161 maintains contact only with the Ala215 and Asp218 of β-propeller. Primarily, Arg of the RGD motif tends to form a salt bridge with Asp218 of β-propeller, while the homo-arginine (HRG) of var-hFn10 with extended alkyl chain (an additional carbon) formed additional interactions with Thr212 and Ala213, besides forming the salt bridge with Asp218. However, the homo-arginine (HRG) of var-hFn10 loses its contact with other β-propeller residues, including Asp150, Phe177, Tyr178, and Ala215. Overall, all RGD ligands maintain interactions consistently with Tyr122, Asn215, and MIDAS Mn^2+^ of the βI domain.

**Fig. 3.**
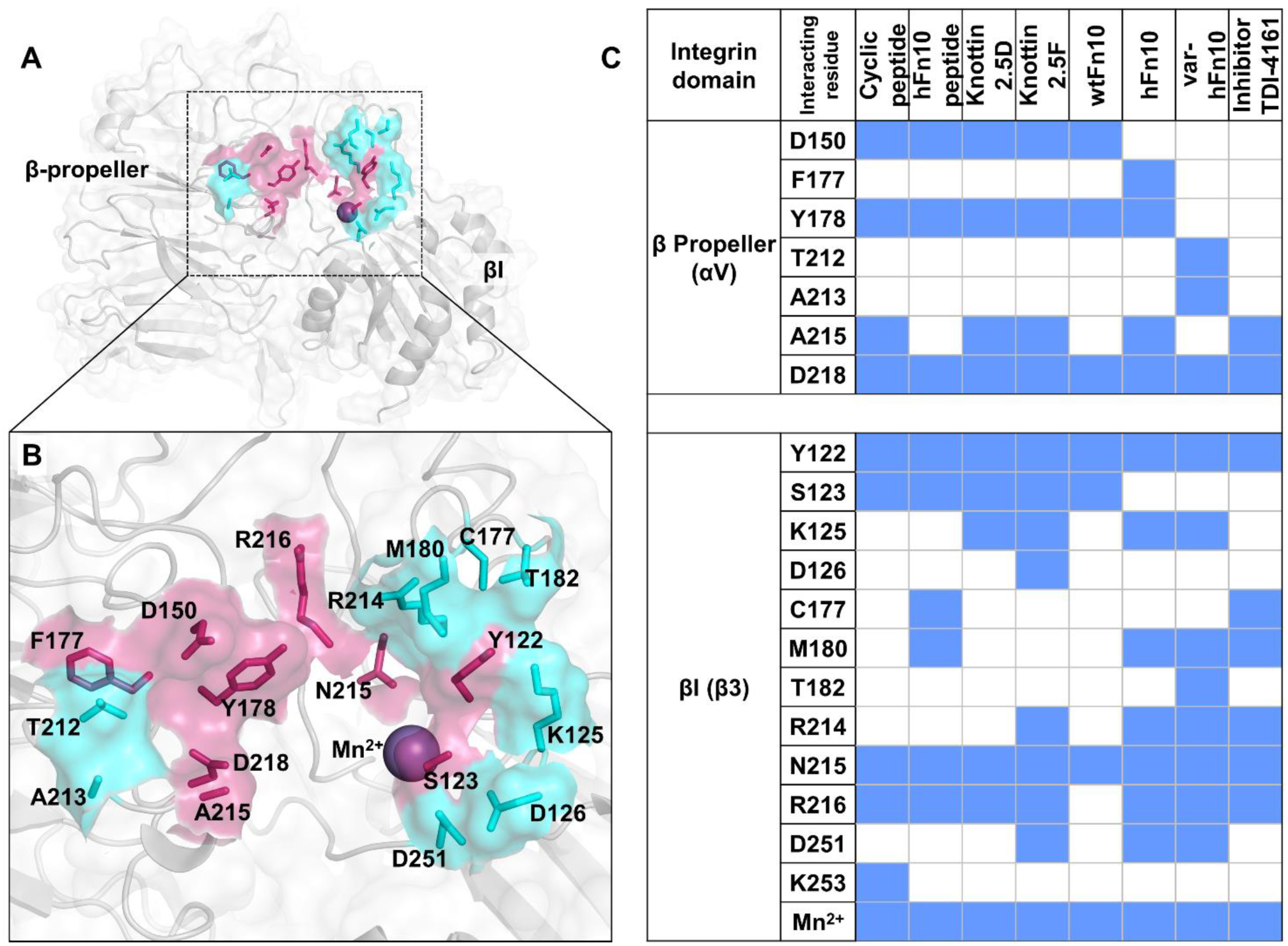
The critical residues of the primary RGD-binding site of integrin αVβ3 and interactions of various ligands. **A-B)** The binding site residues interacting with the RGD motif of the ligands were colored warm pink, and the residues that interact with non-RGD motif residues were highlighted in cyan. Amino acids are represented as sticks and surfaces, and Mn^2+^ as a sphere. Besides residues of αV and β3 chains, the Mn^2+^ ion at the MIDAS site coordinates with Asp of the RGD motif of ligands, and this metal coordination is conserved in all ligand-bound structures. Mn^2+^ is considered as part of the binding site. **C**) Heatmap of the ligand interactions. The presence of an interaction is indicated by blue, while its absence is indicated by white. The heatmap includes all nonbonded interactions between αVβ3 and the RGD ligands. Specifically, interactions with Asp150, Tyr178, Ala215, and Asp218 of the β-propeller domain, Tyr122, Ser123, Asn215, Arg216, and MIDAS Mn^2+^ ion of the βI domain, were found to be conserved among many ligands. With the extended alkyl chain, homo-arginine (HRG) of var-hFn10 did not make contact with Asp150, Phe177, Tyr178, and Ala215. Instead, it interacts with Thr212 and Ala213 of β-propeller. Only antagonists, such as hFn10, TDI-4161, and var-hFn10, were found to interact with one or more of the residues, Cys177, Met180, and Thr182 of SDL in the βI domain.

**Figure 4.**
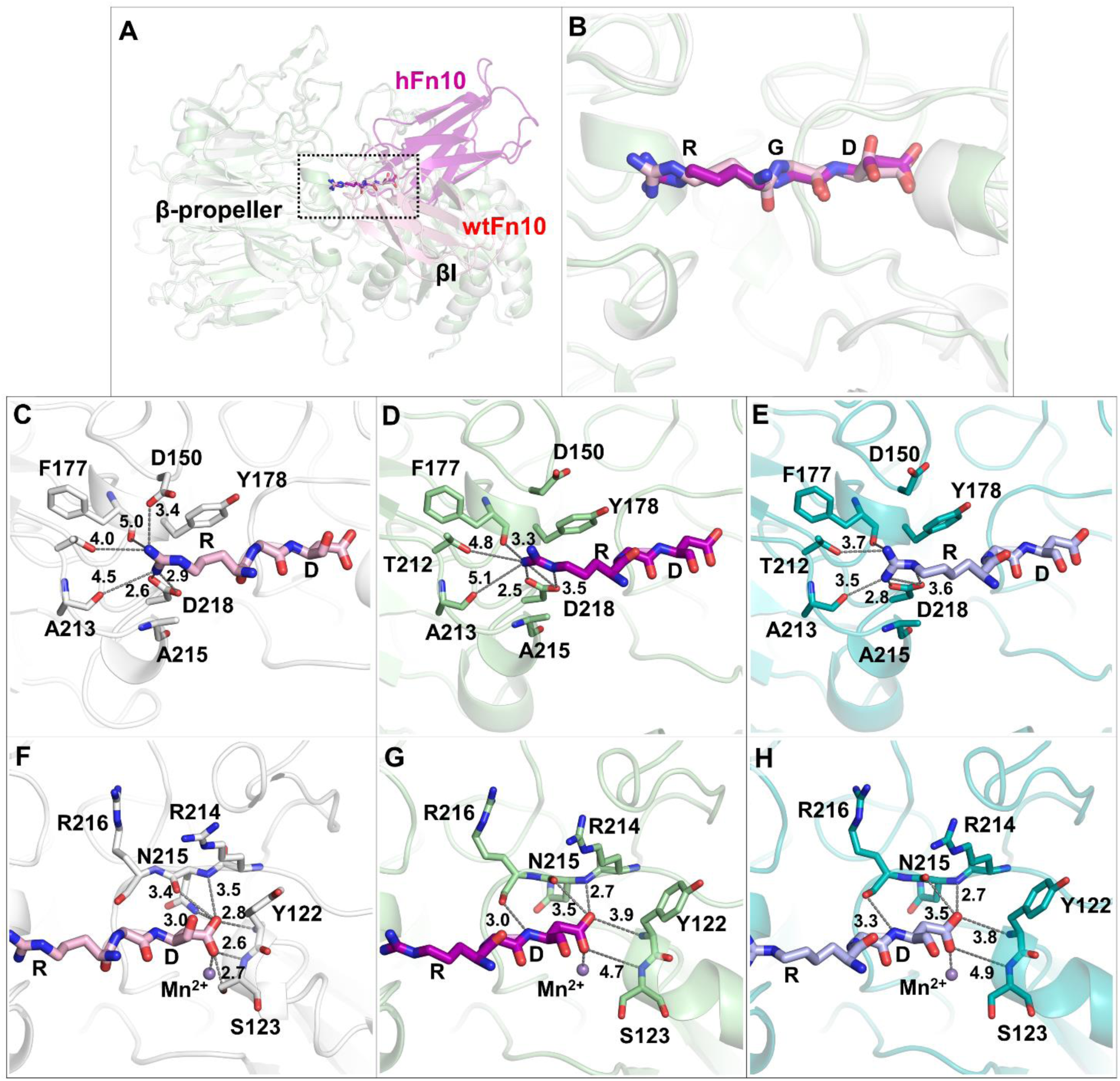
Comparison of RGD motif interactions in agonist and antagonist bound αVβ3 headpiece. **A)** The binding orientations of wtFn10 (light pink color) and hFn10 (magenta) at the RGD binding site of integrin αVβ3. **B)** The orientations of the RGD motif of wtFn10 (light pink color) and hFn10 (magenta) at the site. Despite the differences in the overall orientations of the ligands, their RGD motif residues bind in similar binding modes. **C-E)** The interactions of Arg of the RGD motif in wtFn10 (C), hFn10 (D), and var-hFn10 (E), respectively, show that Arg of hFn10 and var-hFn10 do not interact with D150 in both structures instead forms an H-bond and an electrostatic contact with the backbone carbonyl oxygen of F177 in hFn10 and varFn10 bound structures respectively. The extended alkyl chain-containing homo-arginine of var-hFn10 forms additional contacts with T212 and A213, along with the conserved salt bridge with D218. **F-H)** The interactions of Asp of RGD motif in wtFn10 (F), hFn10 (G), and var-hFn10 (H) show that its interacting distance of Asp with backbone amide nitrogen of Arg214 and the backbone carbonyl oxygen of Arg216 were decreased only in hFn10 (G) yet Asp (RGD) lost contacts with the residues Tyr122, Ser123 in both hFn10 and varFn10 bound structures (G, H). Irrespective of the residue contact differences, the Mn^2+^ ion – Asp (RGD) coordination is conserved in all the RGD ligands.

Specifically, the lack of interaction between the ligands and Ser123 of βI in the hFn10-, inhibitor TDI-4161-, and var-hFn10-bound structures signifies the specificity of the ligands that exhibits pure antagonism. Other than this Ser123-specific interaction, pure antagonists were also observed to interact with Lys125, Cys177, Met180, and Asp251 residues of the βI domain. Likewise, the knottin peptide EETI 2.5F also forms contact with these residues, which could be due to the increased flexibility of the peptide itself[40] (**Fig. 3C**). Though binding is similar to hFn10, var-hFn10 forms an increased number of contacts with the βI domain, and the least number of contacts with the β-propeller domain. This illustrates that the induced binding of var-hFn10 and to accommodate the homo-arginine in the β-propeller domain extends towards the βI domain resulting in an increased number of contacts. The interaction heatmap also reveals that the ligands were able to interact with the SDL residues C177, M180, and T182, and the interactions were specific for the antagonist ligands (**Fig. 3C**).

Despite a significant difference in the overall binding orientations of wtFn10 and hFn10 at the primary binding site, the RGD motif of wtFn10 and the hFn10 binds in a very similar binding mode. However, there are notable differences in their molecular interactions with the site residues (**Fig. 4A and 4B**). The arginine of the RGD motif of the ligands positioned at the narrow groove of the β-propeller domain primarily interacts with the residues Asp150, Phe177, Tyr178, Thr212, Ala213, and Asp218. In the case of wtFn10-bound active structure, the sidechain amino group (NH1) of the arginine residue (of RGD) was positioned near the carboxyl oxygens of Asp218, forming a bidentate salt bridge at distances of 2.6 and 2.9 Å, respectively. The arginine’s charged amino group also formed an H-bond (3.4 Å) with Asp150, while its alkyl chain formed an aryl-alkyl contact with the sidechain aromatic ring of Tyr178 (**Fig. 4C**). In the hFn10-bound inactive structure, the guanidine moiety of the arginine sidechain was rotated, and hence it formed an H-bond (3.3 Å) with the backbone carbonyl group of Phe177 along with the similar bidentate salt bridge with Asp218 at distances 2.5 and 3.5 Å respectively. Unlike wtFn10, H-bond with Asp150 was absent in the hFn10-bound structure due to the slight displacement of the loop as well as the rotation of the Asp150 sidechain away from the arginine. (**Fig. 4D**). Further, interactions of homo-arginine substitution in the var-hFn10-bound structure were analyzed to understand the significance of additional carbon in the arginine’s sidechain that could alter the interactions at the RGD binding site. With the extended chain, the homoarginine moved closer and formed additional electrostatic contacts with Thr212 (3.7 Å) and Ala213 (3.5 Å) residues. While facilitating other contacts through the extended alkyl chain, HRG sidechains were still in the vicinity of Asp218 to maintain electrostatic interactions (**Fig. 4E**). Interestingly, the rotation of the Asp150 sidechain away from the ligands’ arginine was conserved in all the antagonist bound structures.

The difference in the distance between the positive (Arg) and negative (Asp) centers of the peptide, i.e., the distance between R and D in the RGD-motif, has been shown to affect the ligand’s subtype selectivity for the RGD-binding integrins[45]. Remarkably, the additional methyl group in the sidechain of HRG in var-hFn10 offered the ligand a higher affinity for αII_b_β3 as compared to αVβ3. Although the arginine of the RGD motif seems to influence the binding affinity, its absence did not affect the ability of the peptide to activate integrin αII_b_β3[46].

Similarly, Asp of the RGD ligands binds at the cleft formed by the loops (A′-α1, C′-α3, and SDL) of the βI domain and forms a network of polar interactions with the residues Tyr122, Ser123, Arg214, Asn215, Arg216, and Mn^2+^ at the metal ion-induced adhesion site (MIDAS). Coordination with the metal ion is an integral part of ligand binding, which is highly conserved in all the RGD ligand-bound structures. In the wtFn10 bound structure, the carboxyl group of the ligand’s aspartic acid is proximal to α1 helix residues to form H-bonds with backbone NH atoms of Tyr122 (2.8 Å) and Arg214 (3.5 Å), as well as with backbone carbonyl oxygen of Asn215 (3.4 Å) and sidechain NH of Asn215 (3.0 Å). In addition, the carboxyl group of the ligand’s aspartic acid also coordinates with the Mn^2+^ ion and forms two H-bonds with the backbone NH and sidechain OH of Ser123 from α1 helix at distances 2.6 and 2.7 Å, respectively (**Fig. 4F**). In the case of hFn10, the movement of the α1 helix was restrained; thus, the two carboxyl groups did not form any H-bonds with backbone NH atoms of Tyr122 and Ser123 as they remained distantly at 3.9 and 4.7 Å, respectively. Similarly, the backbone carbonyl oxygen atom of Ser123 remained at 6.2 Å from the ligand’s aspartic acid. The H-bonds with NH of Arg214 (2.7 Å) and the carbonyl oxygen of Asn215 (3.5 Å) were maintained in the hFn10-bound structure similar to wtFn10. The sidechain NH of Asn215 has drifted away from the ligand, so its contact with Asp (RGD) was absent in the hFn10 bound structure (**Fig. 4G**). Almost all interactions of Asp of var-hFn10 were similar to that of hFn10 (**Fig. 4H**). As discussed earlier, during the activation of integrin αVβ3, the α1 helix moves inward and closer to the RGD binding site (**Fig. 1B**) so that the α1 residues Tyr122 and Ser123 could form H-bonds with ligand’s Asp residue. In the case of antagonist-bound structures, movement of the α1 helix was restrained, and thus, the residues Tyr122 and Ser123 are distant from the ligands’ Asp in the liganded-inactive conformation.

### Receptor activation accompanies changes in the Mn^2+^ coordination characteristics at MIDAS, ADMIDAS, and SyMBS

It has been known that divalent cations, such as Mg^2+^ and Mn^2+^, can regulate the Integrins’ activation and ligand affinity[47–49]. Comparatively, Mn^2+^ has over ∼40-fold higher affinity for the binding sites than Mg^2+^[48]. In the case of αVβ3, metal ions increase the affinity of integrins towards wtFn10 by ten-fold[47, 50]. The β-propeller and βI domains have several Mn^2+^ ions that are critical structural components of the receptor and were shown to play an imminent role in ligand binding and the receptor activation processes. We analyzed the residue interactions and coordination properties of the Mn^2+^ ions in the unliganded, ligand-bound active, and inactive structures. All the ‘RGD’ ligand-bound structures have three Mn^2+^ ions at the βI domain, coordinating with residues at ADMIDAS (Adjacent to Metal Ion-Dependent Adhesion Site), MIDAS (Metal Ion-Dependent Adhesion Site), and SyMBS/LIMBS (Synergistic Metal-Binding Site/Ligand-Induced Metal-Binding Site) sites (**Fig. 5A**). All the integrin αVβ3 structures contain an Mn^2+^ ion at ADMIDAS, whereas Mn^2+^ ions are observed at MIDAS and SyMBS only in the liganded structures (**Fig. 5A**). The coordination properties of Mn^2+^, such as metal ion occupancy, B-factor, bond valence, coordination geometry, gRMSD, vacancy, and bidentate coordination, are summarized in **Supplementary Table 1** [51]. The coordination angles of Mn^2+^ with various residues and water molecules were summarized in **Supplementary Table 2**. Generally, Mn^2+^ ions can form up to six coordination bonds with ligands and adapt diverse coordination geometries.

**Figure 5.**
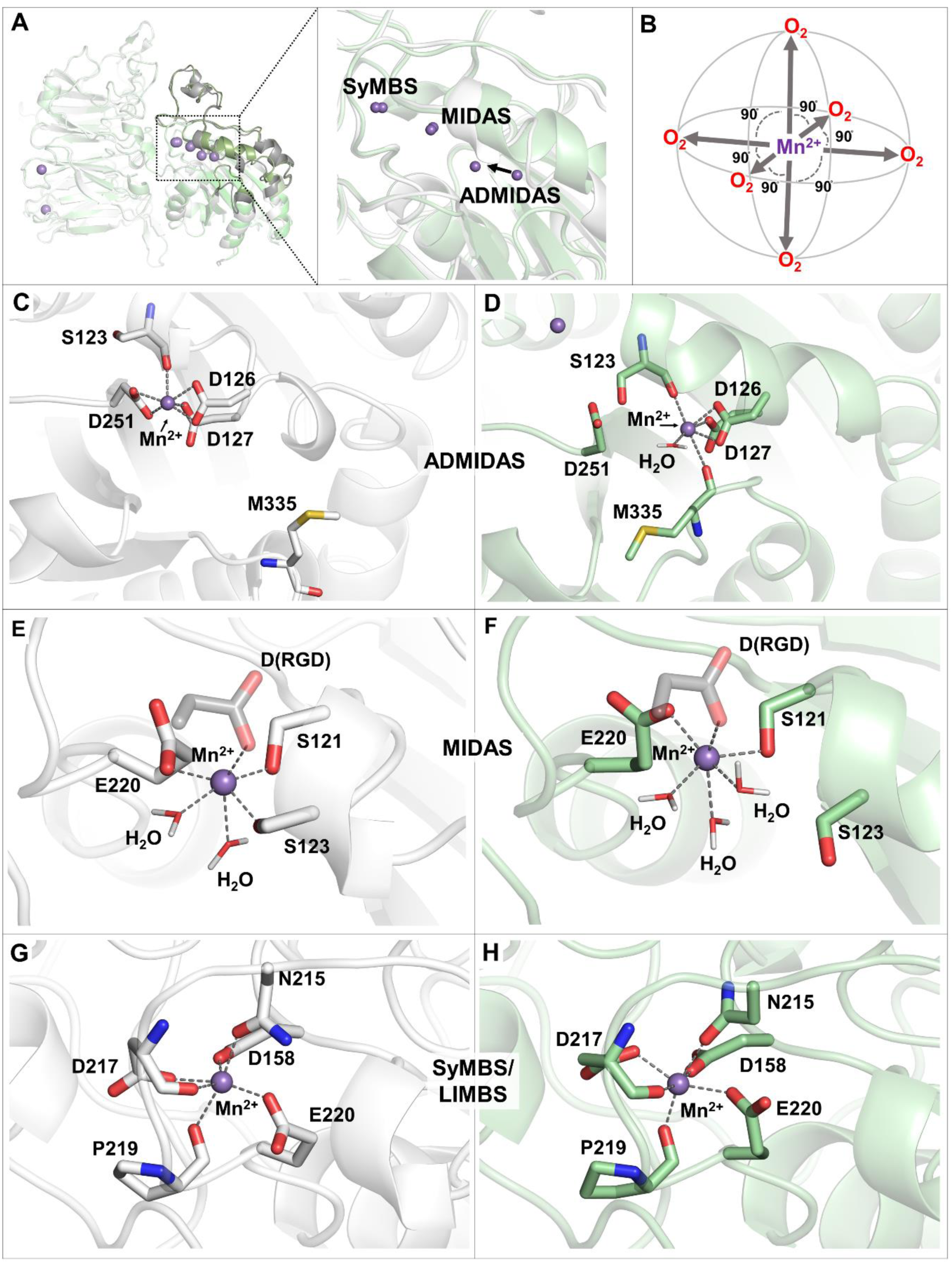
Coordination geometries of metal ions (Mn^2+^) at the RGD binding site. **A)** Integrin αVβ3 has three Mn^2+^ ions at the RGD binding site, occupying ADMIDAS (Adjacent to Metal Ion-Dependent Adhesion Site), MIDAS (Metal Ion-Dependent Adhesion Site), and SyMBS/LIMBS (Synergistic Metal-Binding Site/Ligand-Induced Metal-Binding Site). The coordination properties were compared between wtFn10-(white) and hFn10-bound (green) structures. Only Mn^2+^ ion at ADMIDAS undergoes displacement upon activation, whereas ions at both MIDAS and SyMBS remained in the same positions. **B)** Ideally, Mn^2+^ assumes the octahedral coordination geometry with their ligands at right angles (90 degrees). In αVβ3 structures, all Mn^2+^ ions coordinate with oxygen atoms of the coordinating residues. C-D) In the liganded-active conformation. **C)** At ADMIDAS, Mn^2+^ coordinates with Asp251, Ser123, Asp126, and Asp127. It doesn’t coordinate with Met335 of the <u>F-α7</u> loop, which facilitates the opening of the loop during activation. **D)** In the liganded-inactive conformation, Met335 coordination is retained, and thus the F-α7 loop remains in the inactive conformation. **E)** In the liganded-active structure, Mn^2+^ at MIDAS coordinates with Asp of the RGD ligand, Ser121, Ser123, Glu220, and two water molecules. **F)** In the liganded-inactive conformation, the loss of the Mn^2+^ ion coordination with Ser123 was compensated by coordination with an additional water molecule. **G-H)** The coordination of Mn^2+^ at SYMBS remains the same in both liganded-active and inactive structures.

At ADMIDAS, a difference exists among the residues coordinating Mn^2+^ between the active wtFn10-bound and inactive hFn10-bound structures. In the wtFn10-bound active conformation, the ion coordinates with the backbone carbonyl group of Ser123 and the side chain carboxyl groups of Asp126, Asp127, and Asp251. Among these four residues, Asp126 and Asp251 appear to form bidentate coordination with the metal ion through their carboxyl oxygen atoms and thus contributing to the coordination number of 6 (**Fig. 5C**). In hFn10-bound inactive conformation, Mn^2+^ is coordinated by Ser123, Asp126, Asp127, Met335, and a water molecule. The position of the sidechain of S123 facing Mn^2+^ appears to hinder Asp251 from coordinating with the metal ion.

However, the loss of coordination with Asp251 is compensated by interactions with the backbone carbonyl group of Met335 and a water molecule (**Fig. 5D**). During the receptor activation process, at ADMIDAS, coordination of Mn^2+^ with Met335 was swapped by Asp251 and thus releasing constraints on the F-a7 loop and Met335, allowing the loop to open and move away from the binding site. In the hFn10-bound inactive conformation, the coordination of Met335 with Mn^2+^ seems to prevent the loop opening and, thus, activation. The coordination angles among the interacting residues measured in degrees indicate that most of them deviate significantly from the ideal angle of octahedral geometry, i.e., 90 degrees (**Supplementary Table 2**). Importantly, differences are noticed even among identical residues between the active and inactive conformations.

Mn^2+^ at MIDAS is essential for binding RGD-containing ligands to integrin αVβ3 as it coordinates with the Asp residue (of the RGD motif) of the ligands. In the wtFn10-bound structure, in addition to Asp of ligand, Mn^2+^ coordinates with other βI domain residues, Ser121, Ser123, and Glu220, and two water molecules (**Fig. 5E**). In the hFn10-bound structure, coordination with Ser123 was not observed. However, the absence of S123 coordination is compensated by an interaction with a water molecule to maintain the optimum coordination number (**Fig. 5F**). Though the coordination number was maintained in both liganded-active and inactive structures, the coordination angles are different from the Mn^2+^ ideal geometry (**Supplementary Table 2**). Specifically, the water-mediated Mn^2+^ coordination angles were much higher in the hFn10-bound inactive structure than in the active one. For example, the coordination angles are distinct in the following residue-Mn-residue combinations: 1) E220-Mn-Wat1 (78.5° vs. 104.4° in the active and inactive structures, respectively) and 2) E220-Mn-Wat2 (94.7° vs. 133°). On the other hand, the coordination angles involving E220, Mn^2+^, and Asp of the RDG were 121.9° and 84.8° in the active vs. inactive structures, respectively.

Compared to ADMIDAS and MIDAS, SyMBS is located deeper within the RGD-binding site. The Mn^2+^ ion at SyMBS coordinates with Asp158 of SDL and several C′-α3 loop residues, including Asn215, Asp217, Pro219, and Glu220. Although the coordination properties of Mn^2+^ at SyMBS are similar in general, notable differences in the coordination angles were observed for several residue-Mn^2+^-residue combinations among the active (wtFn10-bound) and inactive (nFn10-bound) conformations (**Fig. 5G and 5H, Supplementary Table 1**). For example, the coordination angles are significantly different among the following residues: 1) D158-Mn-D217 (69.1° vs. 101° in the active and inactive conformations, respectively), 2) D158-Mn-E220 (124.2° vs. 89.6°), and 3) D217-Mn-P219 (58.5 vs. 111.4).

Despite variation in their coordination properties, all three Mn^2+^ ions were observed to be in octahedral geometry in both liganded active and inactive conformations of integrin αVβ3. According to a recent study, over 30% of the protein with Mn^2+^ in the Protein Data Bank have a coordination number of 6 and prefer octahedral and distorted octahedral coordination geometries, followed by 21% of structures with a coordination number of 5, forming square pyramidal or trigonal bipyramidal geometry[51]. Among the three Mn^2+^ ions, only ADMIDAS Mn^2+^ was observed to have a vacancy and bidentate metal coordination in both liganded active and inactive states. The vacancy measures the unoccupied sites in the metal coordination, and the liganded active state was observed to have a value of 33%, which is higher than the optimal ion vacancy threshold (>25%). On the other hand, the liganded-inactive structure had a value of only 16 %. Even though both liganded active and inactive states have the optimum coordination numbers, the differences in the coordination angles make 33% of the coordination site unoccupied in the wtFn10 bound (liganded-active) structure. gRMSD (°) is the deviation of the observed coordination angles from the ideal ones. Only the MIDAS Mn^2+^ of the liganded-inactive state (hFn10) was observed to be an outlier, as its gRMSD (23.6°) was higher than the optimal ion threshold of >21.5°[52].

Among the three Mn^2+^ions, those at ADMIDAS and MIDAS are involved in the ligand binding and activation of the receptor. Changes in the coordination angles indicate the conformational rearrangement of the coordinating residues in the receptor upon binding to wtFn10 and hFn10. In addition, the residues Ser123 and Glu220 appear to coordinate with two Mn^2+^ ions in all the liganded conformations. In the liganded-active conformation, Ser123 coordinates with ADMIDAS Mn^2+^ and MIDAS Mn^2+^ through backbone carbonyl oxygen and the sidechain hydroxyl group, respectively. However, the Ser123 sidechain – MIDAS Mn^2+^ coordination is not observed in liganded-inactive conformation. Glu220 coordinates with SyMBS and MIDAS Mn^2+^ ions through its side chain carboxylic acid group in both liganded-active and inactive conformations. Moreover, the Mn^2+^ coordination at SyMBS seems essential for stabilizing the active site loop conformation.

### The interactions of mannose (glycan) at β-propeller with wtFn10 and hFN10

The analyzed crystal structures of integrin αVβ3 revealed many glycans bound to the protein. Specifically, one of the glycan molecules was bound to residue N266 of the β-propeller domain through N-glycosylation (**Fig. 6**). The glycan, alpha-D-mannopyranose-(1-4)-beta-D-mannopyranoze-(−6)-[alpha-D-mannopyranose-(−1-3)]-beta-D-mannopyranose-(1-4)-2-acemido-2-deoxy-beta-D-glucopyranse-(1-4)-2-acetmido-2-deoxy-beta-D-glucopyranse, was bound in an identical manner in the wtFn10- and hFn10-bound structures (**Fig. 6A**). The glycan made several strong H-bond interactions with the sidechain polar groups of Q214 of β-propeller. However, only in the wtFn10-bound structure, the glycan interacted with the ligand. The interactions include H-bonds with the sidechain -OH of Ser1468 and backbone carbonyl groups of Val1444 and Arg1445[27] (**Fig. 6B**). In contrast, in the hFn10-bound structure, the glycan did not interact with the ligand (**Fig. 6C**). The N266 glycosylation has been shown to be important for ligand binding in other αV-containing integrins, such as αVβ6 and αVβ8. The αVβ6 integrin mediates the cell entry of the Foot-and-mouth disease Virus (FMDV) through RGD motif attachment[53]. The FMDV virus has the propensity for sulfated-sugar binding[53], which further extends the functional importance of Asn266 glycosylation in the β-propeller of the αV subunit. There is also an N-glycan found at the corresponding position on integrin α5β1 at Asn275 of α5. The crystal structure of α5β1 with the 7-10^th^ domains of fibronectin (Fn7-10) shows that the glycan at the Asn275 of α5β1 (equivalent to Asn266 of αVβ3) extends towards the interface between Fn9 and Fn10 (**PDB ID: 7NWL**)[54]. Although the glycan only forms interactions with the Fn9 domain of fibronectin. The N275A mutation reduced the binding affinity of fibronectin by ∼60%, reinforcing the importance of glycosylation at this position[55]. Additionally, the N275Q mutation impaired the α5β1-mediated cell adhesion[56]. Furthermore, the glycan-fibronectin interactions differ among these integrin subtypes due to the bound orientation of the fibronectin. We compared the binding orientation of the tenth domain of fibronectin in αVβ3 and α5β1. For this purpose, we measured the angle formed by the center-of-mass (COM) of the fibronectin Fn10 domains by keeping the RGD motif as the center of the angle. In α5β1, fibronectin moved ∼15° towards the βI domain of β1 than in αVβ3. Also, in αVβ3, the high-affinity fibronectin (hFn10) moved around ∼40.4° towards β3 than in wtFn10. These experimental and structural pieces of evidence suggest that the glycan interactions may differ among integrin subtypes and validate the significance of N-linked glycan at Asn266 for ligand binding and affinity.

**Figure 6.**
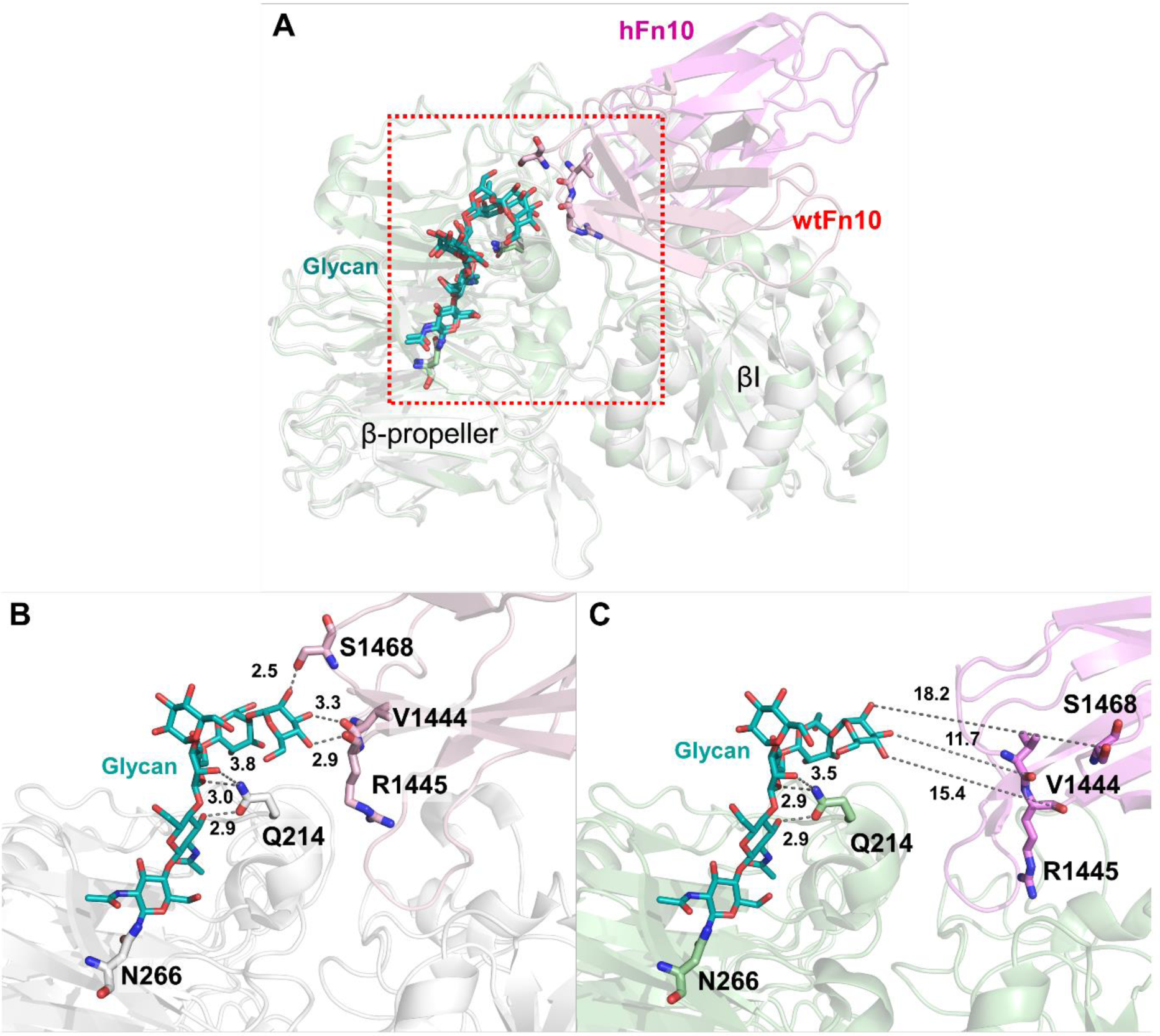
Glycan interacts with wild-type fibronectin. **A)** Comparison of the bound orientation of the glycan on the wtFn10 (light pink) and hFn10 (violet) bound active (white) and inactive (green) integrin structures shows that the glycan is shown as teal-colored sticks and Asn266 in the β-propeller of αV is glycosylated in both active and inactive conformations of the integrin. **B)** Moreover, the glycan moiety is bound in a similar orientation, and the sugar moiety interacts with Glu214 of the β-propeller and Val1444, Arg1445, and Ser1468 of wtFn10. **C)** In the inactive structure, hFn10 lost contact with the glycan.

### Antagonists engage in distinct interactions as compared to wtFn10

Despite engaging in similar interactions through their RGD motif at the primary binding site, the binding orientation of wtFn10 is markedly distinct from that of hfFn10 and var-hFn10 (**Fig. 7A and 7B**). This difference in the binding orientation facilitates hFn10 and var-hFn10 to form additional contacts with the βI domain residues in the vicinity of the RGD-binding site. In contrast, no significant interactions were found with wtFn10 other than the primary RGD motif interactions (**Supplementary** Figure 1). In hfFn10, Trp1496 forms a π-π interaction with Tyr122 of α1 helix spanning the βI domain. This Trp1496–Tyr122 interaction confines the position of Tyr122, enabling additional interactions formed by hFn10 in the inactive receptor conformation. Also, this π-π stacking interaction appears to arrest the inward movement of the α1 helix towards site-I, one of the essential activation signatures of the integrin αVβ3 headpiece. The hindrance in the inward movement of the α1 helix disrupts the ability of Asp (of the ligand RGD-motif) to form H-bonds with the backbone atoms of both Tyr122 and Ser123 of the α1 helix (**Fig. 4G**). While the backbone (NH) and sidechain (OG) atoms of Ser123 are positioned distantly at 4.7 Å and 6.2 Å, respectively, in the hfFn10 bound structure, the corresponding distances are 2.6 Å and 2.7 A in the wtFn10-bound active structure (**Fig. 4F and 4G**).

**Figure 7.**
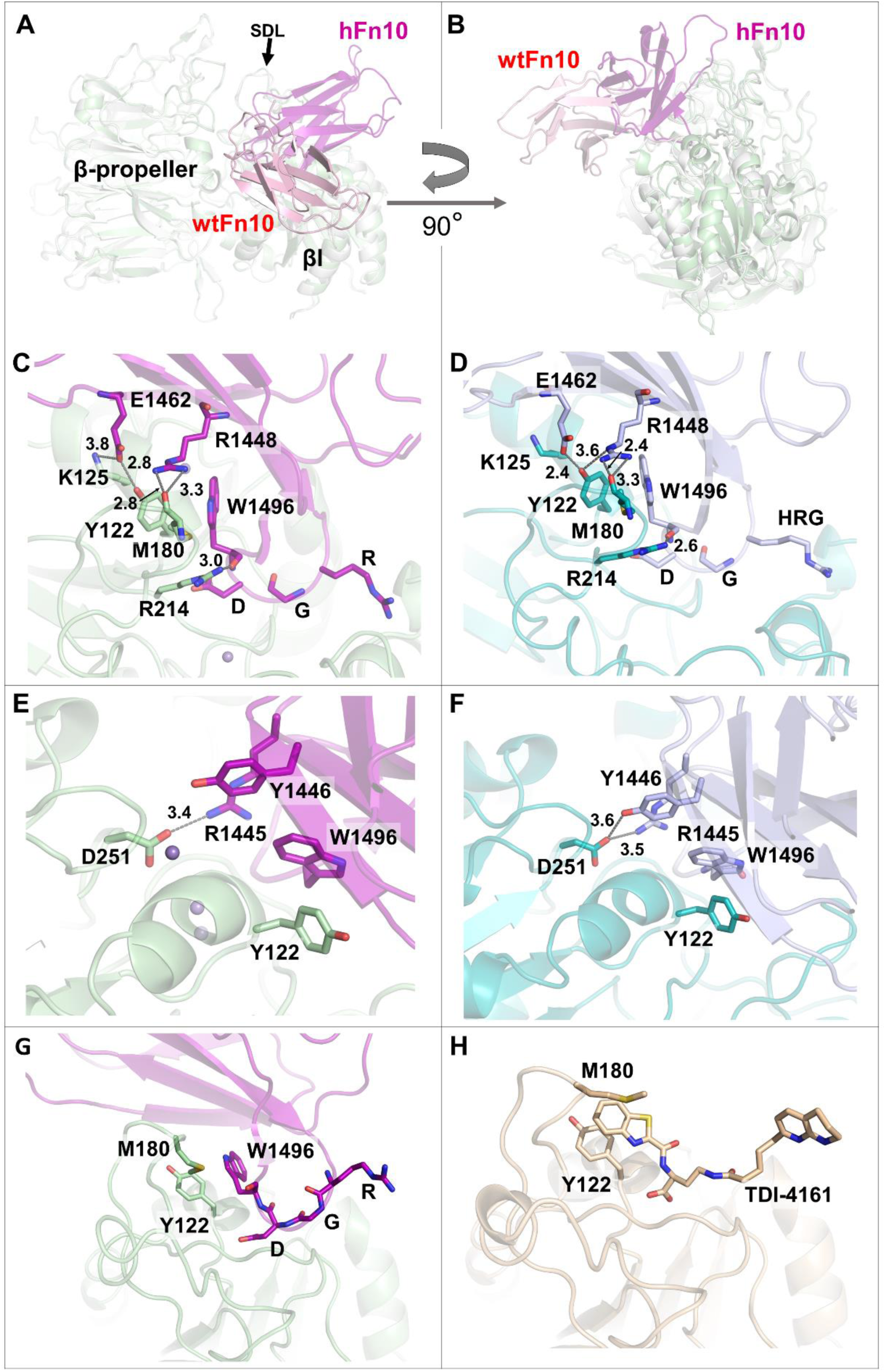
wtFn10 and hFn10 bind to integrin αVβ3 in distinct orientations, and antagonists form specific interactions with SDL. **A-B)** The distinct binding orientations of wtFn10 (light pink) and hFn10 (magenta) are shown in the top and side views, respectively. WtFn10 is oriented towards the α7 helix of the βI domain, whereas htFn10 is oriented towards the α1 helix and the SDL loop. The difference in the binding orientation facilitates hFn10 to form many additional contacts with the βI domain. Var-hFn10 binds in orientation like hFn10 and engages in similar interactions. **C)** In hFn10, in addition to the π-π interaction of ligand Trp1496 with Tyr122, residues Arg1448 and Glu1462 also form electrostatic interactions with the βI domain. Specifically, the guanidine sidechain of Arg1448 forms an H-bond with the backbone carbonyl group of Met180, and Glu1462 forms electrostatic contacts with sidechains of Tyr122 and Lys125 of the βI domain. **D)** In var-hFn10, Arg1448 forms H-bonds with the backbone of Met180 and with the sidechain of Tyr122. Specifically, Glu1462 forms electrostatic contact with Tyr122, but its contact with Lys125 is lost. E) Arg1445 of hFn10 forms a salt bridge with Asp251 of the ADMIDAS site. **F)** In var-hFn10, residues Arg1445 and Tyr1446 form electrostatic interactions with Asp251. **G)** In the hFn10-bound structure, Trp1496 is positioned to form a π-π interaction with Tyr122 and does not make any contact with Met180. **H)** In addition to a π-π interaction with Tyr122, the small molecular inhibitor TDI-4161 forms an aryl-alkyl interaction with the sidechain of Met180.

In addition to Trp1496, Arg1445, Tyr1446, Arg1448, and Glu1462 of hFn10 also interact with the βI domain. In the hfFn10-bound structure, Glu1462 forms an H-bond (2.8 Å) with the sidechain hydroxyl group of Tyr122 and electrostatic interactions with the sidechain amino group of Lys125 (3.8 Å). Besides, the nitrogen atoms of the sidechain guanidinium group of Arg1448 are positioned at distances of 2.8 and 3.3 Å and form H-bonds with the backbone carbonyl oxygen of Met180 from the SDL loop (**Fig. 7C**). The positively charged amino group of Arg1445 of hfFn10 forms a salt-bridge with the negatively charged carboxylic group of Asp251 from the β sheet that is positioned directly below the α1 helix (**Fig. 7E**). Like hFn10, var-hFn10 has Ser1496Trp mutation, and forms interactions similar to that of hFn10 but with minimal changes in the interaction pattern (**Fig. 7D and 7F**). In the var-hFn10-bound structure, Glu1462 forms an H-bond with Tyr122, whereas Arg1448 forms similar H-bonds with the backbone carbonyl oxygen of Met180. In addition, the Asp251 of the βI domain forms a salt bridge and electrostatic interactions with Arg1445 and Tyr1446 of var-hFn10. Besides var-hFn10, αVβ3 selective small molecule inhibitor TDI-4161 was reported to have a similar inhibition profile as hfFn10. Since TDI-4161 was designed to mimic the ideal interactions of the **RGDW** motif of hfFn10, TDI-4161 forms identical interactions at the RGD binding site and exhibits similar conformational arrest of the headpiece, resulting in the inhibition of the receptor. The benzothiazole moiety from TDI-4161 is harbored between the SDL loop and forms a parallel π-π stacking interaction with the Tyr122 sidechain (**Fig. 7H**). Although Trp1496 of hFn10 and varFn10 engages in similar π-π stacking interactions with Tyr122, the relative distance and stacking geometry seem different. Specifically, the distance between the phenyl ring of Tyr122 and the center of the indole ring of Trp1496 of hFn10 and varFn10 are 5 Å and 4.7 Å, respectively. Similarly, the angle between the two interacting aromatic ring planes for hFn10 and varFn10 are 43° and 55°, respectively (**Supplementary** Figure 2). In the case of TDI-416, the aromatic rings of Tyr122 and TDI-4161 were stacked more closely (at a distance of 4 Å) and parallelly (at an angle of 11°). The macromolecule inhibitors, hFn10 and var-hFn10, form more contact with the βI domain than TDI-4161. Amongst them, the H-bonds with the Met180 backbone carbonyl oxygen are one of the consistent interactions observed in both hFn10 and var-hFn10 bound structures. In contrast, an aryl-alkyl interaction with the Met180 sidechain is observed for TDI-4161, which was absent in both hFn10 and var-hF10 (**Fig. 7F**).

### Site II mediated activation of αVβ3

In addition to many RGD motif-containing ligands, αVβ3 integrin is also functionally regulated by non-RGD mediators such as chemokine fractalkine, pro-inflammatory secreted phospholipase A2 type IIA (sPLA-IIA), CXCL12, CD40L, and oxysterols such as 25-hydroxycholesterol (25HC) and 24(S)-hydroxycholesterol (24HC)[17, 57]. Though substantial experimental evidence is available for the integrin activation by these non-RGD ligands, the structural basis for the activation mechanism is yet to be elucidated. Prediction of the binding orientations of these non-RGD ligands based on molecular docking and molecular dynamics simulations revealed that the non-RGD ligands might regulate integrin activation allosterically through binding at the region distinct from the primary RGD binding site. The non-RGD ligands are predicted to bind at a site located at the interface of the β-propeller and βI domains and distant (∼ 6Å) from the primary binding site. The site encompasses the residues, including Glu15, Tyr18, Lys42, Asn44, and Gly49. Ile50, Val51, Glu52, Asn77, Ser90, His91, Trp93, His113, Arg122, Ala397, Arg398, Ser399 and Met400 of the β-propeller domain and Val161, Ser162, Ala263, Gly264, Ile265, Gln267, Asp270, Gln272, Cys273, His274, Val275, Gly276, Ser277, Asp278, Asn279, His280, Tyr281, Ser282, Ala283, Ser284, Thr285, Thr286 and Met287 of the βI domain, respectively (**Supplementary** Figure 3). Although the ligands were observed to be in contact with αVβ3 via site II, the exact mechanism by which the ligands induce conformational changes and lead to receptor activation is yet unclear. In our recent study, we showed that during viral infections, macrophages produce 25HC, an oxygenated metabolite of cholesterol, which can bind directly to αVβ3 and α5β1 integrins with nanomolar affinity and induce pro-inflammatory responses. Surprisingly, 25HC is the first endogenous lipid molecule reported to be a regulator of integrin activation. Our in vitro and computational studies revealed that 25HC directly binds to site II of integrins αVβ3 and α5β1 and increases the production of pro-inflammatory mediators such as IL-6 and TNF by activating the integrin-FAK pathway. Molecular docking and molecular dynamics simulation studies showed that 3-OH and 25-OH of 25HC form stable interactions with Ser162 and Ser399 residues of site II [17]. The binding of 25HC at site II has produced significant conformational changes in SDL. The observation that unlike wtFn10, antagonists such as hFn10, var-Fn10, and TD1-4161 directly interact with the SDL residues and produce differential functional outcomes suggest that the primary RGD-binding site may be allosterically modulated via SDL by targeting site II.

### SDL loop conformation and H-bond networks

The specificity-determining loop (SDL) is made up of a ∼30 amino acid segment (residues 158 to 190) at the βI domain (**Fig. 8**). As the name suggests, SDL plays a crucial role in determining the specificity of ligand recognition by integrin αVβ3. Even though part of αVβ3 SDL is positioned near the β-propeller, neither it is significantly influenced by the β-propeller nor involved in the dimer formation but undergoes substantial alteration in the interactions when it binds with ligands [58]. As seen earlier in Figure 7, residues from SDL are in direct contact with the antagonists, hFn10 and var-hFn10, and the small-molecule inhibitor, TDI-4161 (**Fig. 7H**). Also, the αVβ3-specific monoclonal antibody, LM609, directly binds to the SDL residues and sterically hinders the binding of other extracellular matrix ligands (**Fig. 2**). The secondary structure of SDL appears to be maintained irrespective of the nature of the bound ligands and the activation state of the receptor. The short knot-like conformation of SDL is stabilized by a disulfide bond (between C177 and C184), and interactions of the backbone and sidechain atoms of the SDL residues with the residues located adjacent to the loop in the headpiece and with the residues from the binding partners. Several differences were observed in the inter-residue interactions within SDL in the agonist-(wtFn10) and antagonist-(hFn10) bound receptor structures.

**Figure 8.**
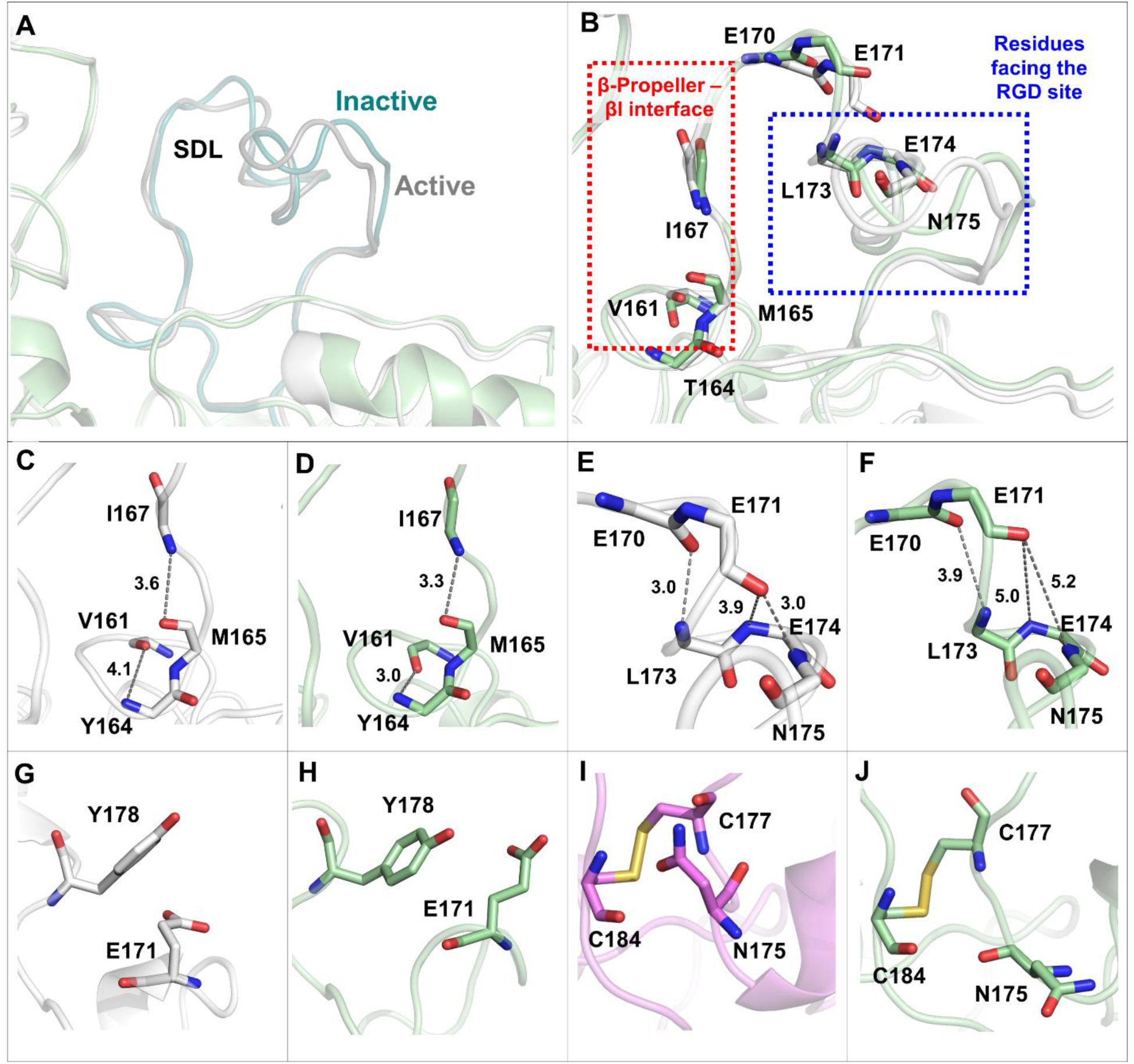
Conformation and altered interactions within the SDL loop. **A)** Superimposition of wtFn10 (white) and hFn10 (pale green) bound structures. SDL loop was highlighted with respective darker colors in liganded-active (teal) and liganded-inactive (grey) conformations. **B)** The backbone atoms of the residues exhibited differences in the interactions and were shown as sticks. **C-D)** The backbone interactions of residue pairs Ile167-Met165 and Val161-Tyr164 at SDL in the wtFn10-(white) and hFn10-(pale green) bound structures, respectively. **E-F)** The backbone interactions of residue pair Glu170-Leu173 and Glu171-Glu174-Asn175 in the wtFn10-(white) and hFn10 (pale green)-bound structures, respectively. The H-bonds between the pairs Glu171-Asn175 and Glu170-Leu173 and electrostatic contacts between Glu171-Glu174 were lost entirely in the hFn10-bound structure. **G-H)** An anion-π contact was observed between the sidechain of Tyr178 and the carboxyl oxygen of Glu171 in the wtFn10-bound structure (white), which was absent in the hFn10-bound structure (pale green). **I-J)** The side chain of Asn175 is positioned closer to the Cys177-Cys184 disulfide bridge in the unliganded (purple) structures, but it is positioned away from the bridge in the hFn10 bound structure.

In the wtFn10-bound structure, at the ligand-binding interface, the backbone electronegative atoms of the residue pair Val161-Tyr164 and Met165-Ile167 form electrostatic contacts with distances of 4.1 Å and 3.6 Å, respectively (**Fig. 8C**). However, in the hFn10-bound structure, the distances were decreased to 3.0 Å and 3.3 Å, indicating stronger contact in the presence of an antagonist (**Fig. 8D**). Additionally, in the wtFn10 bound structure, the SDL segment facing the RGD binding site forms H-bonds between the backbone atoms of Glu170 and Leu173 and Glu171 and Asn175 residues at similar distances (∼ 3.0 Å). In addition, the backbone atoms of Glu171 and Glu174 appear to make electrostatic contacts at a distance of 3.9 Å (**Fig. 8E**). In contrast, in the hFn10-bound structure, the distances between these H-bond pairs Glu170-Leu173 and Glu171-Asn175 were increased to 3.9 and 5.2 Å, respectively. Furthermore, the electrostatic contact between Glu171-Glu174 was absent in the hFn10-bound inactive structure (**Fig. 8F**).

An anion-π interaction between the aromatic ring of Tyr178 and the sidechain carboxyl group of Glu171 was observed in wtFn10 (**Fig. 8G**). However, in the hFn10-bound structure, the orientation of the Glu171 sidechain is significantly altered such that the carboxyl group is engaged in an H-bond interaction with the sidechain -OH group of Tyr178 (**Fig. 8H**). The disulfide bond between Cys177 and Cys184 in the ligand-binding interface of SDL remained intact in all the studied structures. Only in the unliganded structures, the sidechain of Asn175 is positioned near the disulfide bridge, engaging in polar interactions with the backbone carbonyl and amino group of Cys177 (**Fig. 8I**). In contrast, the sidechain appears to move away entirely in the opposite direction in all the ligand-bound structures (**Fig. 8J**).

Overall, in the liganded-active conformation, the inter-residue backbone H-bond distances in SDL appear to be decreased at the β-propeller–βI interface region, whereas the distances were increased between residues facing the RGD binding site (**Fig. 8B**). Thus, in the liganded-active conformation, the SDL segment facing the RGD ligands has also slightly moved towards the β-propeller domain along with the α1 helix. These subtle differences illustrate that the SDL residues undergo conformational changes upon ligand binding and subsequent receptor activation.

### Interactions of SDL with the β-propeller domain

The SDL residues do not have any direct contact with the β-propeller domain in the unliganded structures. However, SDL moved closer to β-propeller upon binding to wtFn10 and moved even closer while binding to hFn10, thereby forming contacts with its residues. Primarily, two serine residues of SDL Ser162 and Ser168 interact with His113, Asp121, and Arg122 of β-propeller (**Fig. 9A**). In the unliganded structure, the sidechain hydroxyl group of Ser168 is 10.1 Å and 7.0 Å away from the sidechain carboxyl group of Asp121 and backbone NH group of Arg122, respectively. These distances decreased to 6.6 Å and 6.2 Å, respectively, in the wtFn10-bound structure. Interestingly, in the hFn10-bound structure, β-propeller and βI domains moved further closer to each other thereby, the sidechain of Ser168 projected towards the β-propeller forming H-bonds with the sidechain carboxyl group of Asp121 and backbone NH group of Arg122 with shorter distances of 3.0 Å and 2.7 Å, respectively than in the unliganded and wtFn10-bound structures (**Fig. 9B**). Similarly, H-bonds were observed between the Ser162 sidechain -OH group and the backbone carbonyl oxygen of Ala263 in the unliganded, wtFn10-, and αV-specific antibody-bound structures with the bond distance of 3.5 Å, 3.3 Å, and 3.0 Å, respectively. However, in the hFn10-bound structure (pale green), the Ser162 sidechain -OH group forms an H-bond (3.5 Å) with the imidazole nitrogen (Nε) of His113 from the β-propeller domain. As Ser162 moved towards the β-propeller domain, the distance between the Ala263 carbonyl group and Ser163 sidechain increased significantly to 5.8 Å (**Fig. 9C**). Interestingly, none of these SDL-β-propeller interactions were observed in the αV-specific monoclonal antibody-integrin complex.

**Fig. 9.**
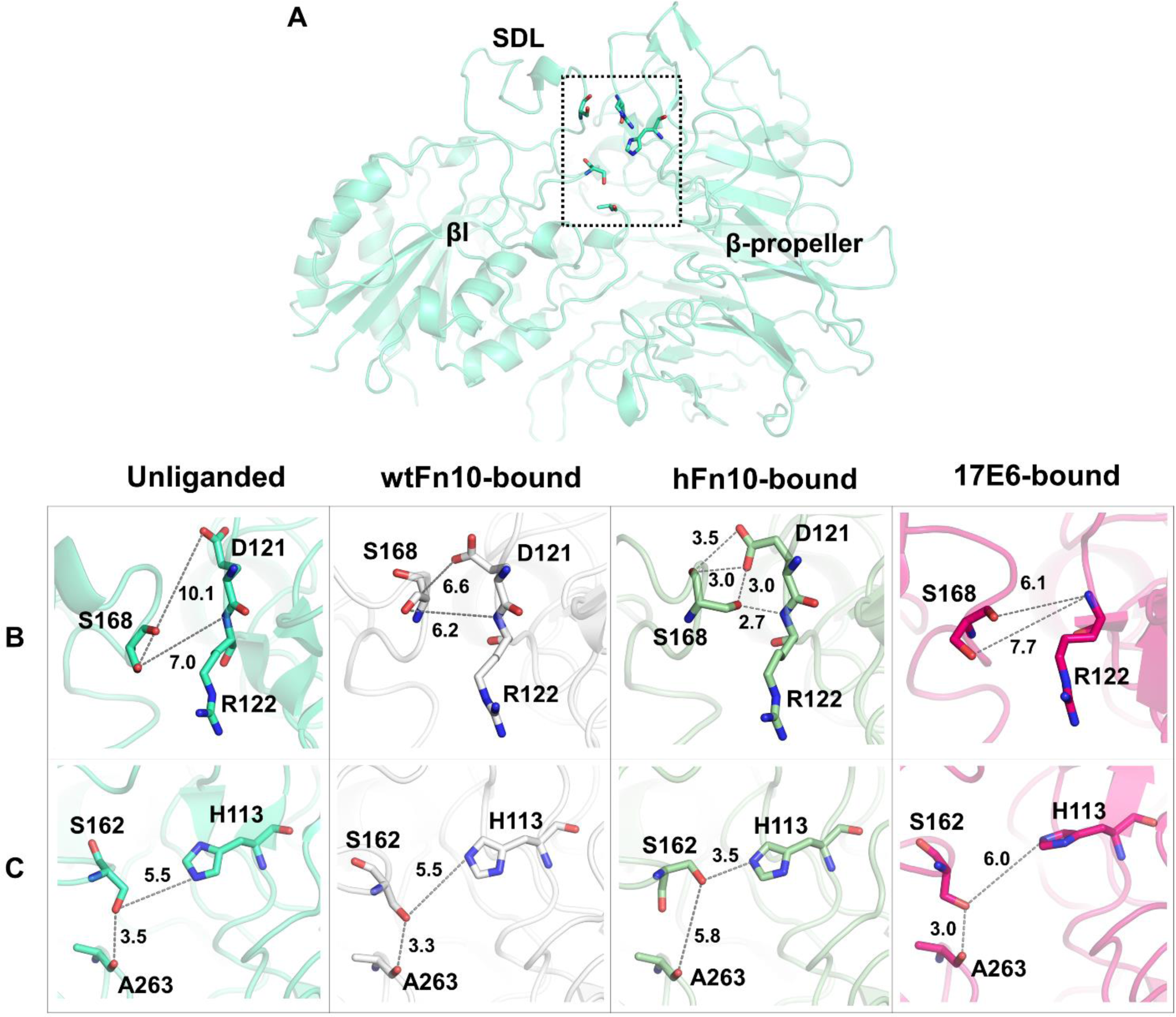
Molecular interactions of SDL with the β-propeller domain. **A)** The interacting residues from SDL and the β-propeller domain are shown in boxes. Primarily, S162 and S168 of SDL interact with H113, D121, and R122 of β-propeller. **B)** In the hFn10 bound structure (pale green), SDL’s Ser168 forms H-bond interactions with the sidechain and backbone atoms of the Asp121 and Arg122 residues of the β-propeller domain, respectively. However, these residues are far apart in the unliganded (cyan), wtFn10 (white)-, and αV-specific antibody bound (magenta) structures and do not form any such interactions observed in the hFn10-bound structure. **C)** Ser162 of SDL does not make any contact with β-propeller in the unliganded (cyan), wtFn10-(white), and αV-specific antibody-bound (magenta) structures. However, in these structures, the sidechain of Ser162 forms an H-bond with the backbone carbonyl group of Ala263. In the hFn10 bound structure (pale green), the backbone carbonyl of S162 moved towards the β-propeller and formed an H-bond with His113 while oriented away from Ala263. Ser162 and Ala263 residues were part of the allosteric site (site II).

### SDL serves as an interface between the primary RGD binding site (site I) and the allosteric site (site II)

Ligands of integrin αVβ3 form contacts with the SDL residues either directly or indirectly through the layer of primary RGD binding site residues. Unlike wtFn10, which lacks any direct contacts, hFn10 forms several contacts with SDL (**Fig. 7**). In the absence of direct contact, the interactions of the RGD motif with the primary binding site residues and differences in their interactions with SDL residues via secondary layers of residues in both wtFn10- and hFn10-bound structures were analyzed. In the wtFn10-bound structure, Asp179 forms a salt bridge with the Arg214 sidechain, which is highly conserved in all the analyzed crystal structures except for the αV-specific antibody bound structure. In addition, the Arg214 sidechain forms electrostatic contacts with the backbone carbonyl groups of Pro176 (3.9 Å) and Asp179 (3.6 Å) and forms H-bonds with the sidechain phenolic -OH group of Tyr166 (2.8 and 3.1 Å) (**Fig. 10A**). As the Asp179 sidechain moves closer to Arg214 in the hFn10-bound conformation, it mediates the formation of a salt bridge with the guanidium group of Arg214, which leads to tighter and rigid association with the SDL loop (**Fig. 10B**).

**Figure 10.**
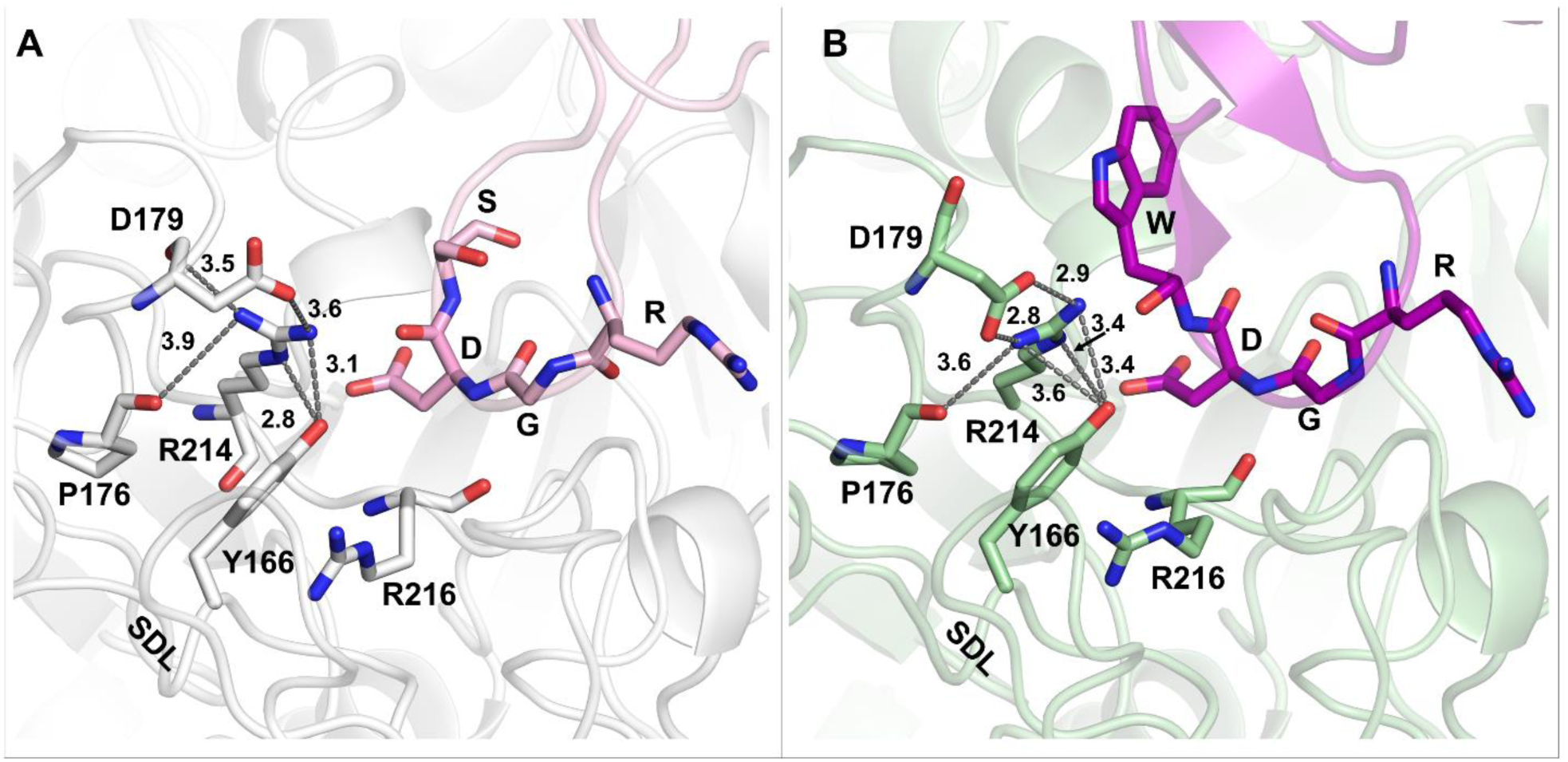
Ligand-induced interactions between the RGD binding site and the SDL residues. Comparison of the RGD interactions in wtFn10 (light pink) and hFn10 (magenta) bound structures shows that the interaction distances were decreased in the hFn10 bound structure. The tighter connection with hFn10 in the RGD binding site induces changes in the sidechain orientations of the active site residues that interact with the SDL. **A)** In the wtFn10 (light pink)-bound αVβ3 (white) structure, Arg214 from the RGD binding site forms an electrostatic contact with Pro176 and Asp179 of SDL and forms H-bonds with the Tyr166 sidechain. The Arg216 sidechain forms a cation–π interaction with Tyr166, and the Arg214 sidechain Nε forms an electrostatic contact with the Tyr166 sidechain. **B)** In the hFN10 (magenta) bound αVβ3 (pale green) structure, the orientation of the Arg214 sidechain was slightly altered, and the guanidino group of the sidechain formed a salt bridge with the Asp179 sidechain carboxyl group. On the other side, the Arg214 sidechain group engaged in a weaker H-bond with the sidechain hydroxyl group of Tyr166. Due to the change in the orientation of the Arg214 sidechain, Tyr166 of SDL forms cation-π interactions with the sidechain guanidium groups of both Arg214 and Arg216, where the phenyl ring of Tyr166 is sandwiched between the two arginine sidechains.

In the hFn10-bound structure, Asp of RGD forms H-bonds with both Arg214 (NH) and Arg216 (CO) through the sidechain carboxylic group and backbone -NH group, respectively with shorter bond distances and led to the changes in the sidechain orientation of both the arginine residues (**Fig. 4G and 4H**). These changes favor Arg214 to form H-bonds with Asp179 and weaker H-bonds with Tyr166 (**Fig. 10B**). Moreover, in the hFn10-bound structure, the sidechains of Arg214 and Arg216 moved further closer towards the SDL and cause the Tyr166 of SDL to be sandwiched between these arginine residues by π-cation interactions (Fig. 10B). On the other hand, in the unliganded and wtFn10-bound structures, only Arg216 forms a cation-π interaction with the aromatic ring of Tyr166 (**Fig. 10A**). These structural analyses of the wtFn10-(active) and hFn10-bound (inactive) αVβ3 structures illustrate that the subtle differences in the interactions of binding site residues with the RGD-ligands affect the contacts between SDL residues and RGD binding site residues.

In the wtFn10 bound structure, the backbone carbonyl oxygen of the Arg216 made an electrostatic contact (3.6 Å) with the backbone -NH of Asp of the RGD ligand. The sidechain guanidinium group of Arg216 was further connected with the backbone carbonyl oxygen of Met165 through an H-bond (3.5 Å). In addition, the backbone -NH group of Met165 made H-bonds with the backbone carbonyl groups of Val161 and Ser162 with distances of 3.1 Å and 2.7 Å, respectively. Apparently, Met165 acts as a linker that connects the RGD binding site residue Arg216 with site II residues Val161 and Ser162 (**Fig. 11A**). In the case of the hFn10-bound structure, it appears that a subtle change in the orientation of the Arg216 sidechain resulted in a relatively stronger H bond (Arg216 sidechain - Met165 backbone) with a distance of 3.3 Å. While Arg216 - Met165 and Met165 – Val161 interactions were conserved in both liganded-active and inactive conformations, the Met165 - Ser162 H-bond was lost in all the inactive conformations in which the receptor is bound to antagonists such as hFn10, var-hFn10, and the small-molecule inhibitor TDI-4161. For example, in the hFn10-bound structure, the Ser162 backbone flipped away from the Met165 backbone NH group, with a distance of 5.7 Å, thereby losing the H-bond (**Fig. 11B**). As mentioned earlier, the flipped Ser162 residue favors the positioning of its sidechain towards β-propeller in the hFn10-bound inactive conformation (**Fig. 9C**). Additionally, Asn215 at the RGD binding site forms weaker H-bonds via its backbone and sidechain amide nitrogen atoms with the carboxylic group of Asp of the RGD ligand (**Fig. 4F**). Also, Asn215’s sidechain carbonyl oxygen forms an H-bond with the backbone carbonyl oxygen of Tyr164. Similar to Met165, Tyr164 also acts as a linker that connects Asn215 with Val161 of site II (**Fig. 8C**). In the antagonist-bound structures, the sidechain amino group of Asn215 flipped and positioned towards the Tyr164 to form an H-bond with the backbone carbonyl oxygen of Tyr164. Despite flipping, the Asn215 -Tyr164 and Tyr164-Val161 contacts were conserved in unliganded and ligand-bound conformations. However, these electrostatic interactions remain weaker in wtFn10-bound conformation as compared to the antagonist-bound inactive conformations. Moreover, the Asp (RGD)-Asn215-Tyr164 H-bond network becomes stronger with shorter bond distances in the hFn10-bound conformation. For example, the contact distance between Tyr164 and Val161 in the hFn10-bound conformation (3 Å) is much smaller than in the wtFN10-bound one (4.1 Å) (**Fig. 8D**).

**Figure 11.**
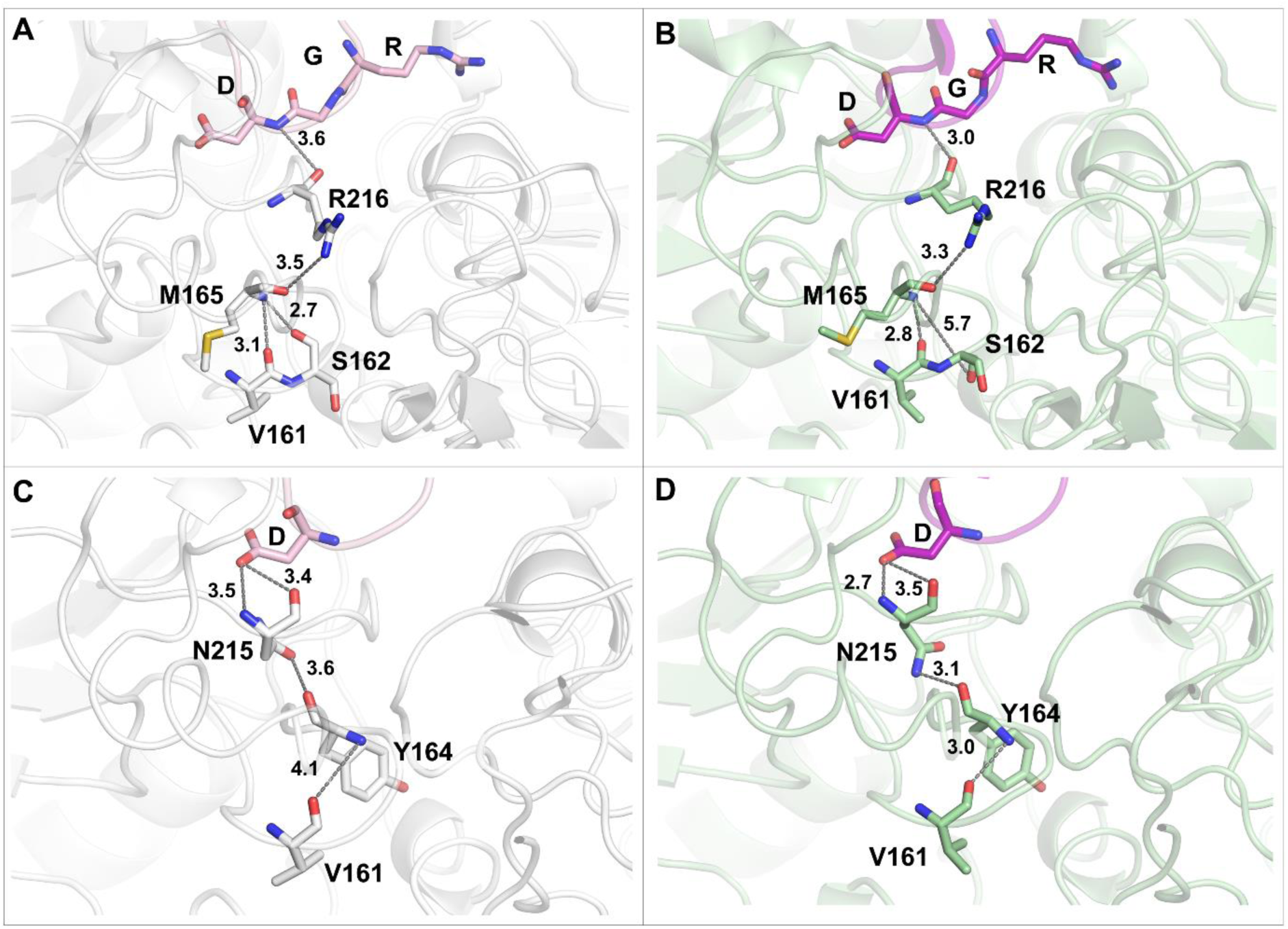
SDL may serve as an interface between the primary RGD binding site and site II. The wtFn10-(active) and hFn10-bound (inactive) conformations of integrin αVβ3 are shown in secondary structure representations in white and pale green, respectively. The ligands wtFn10 and hFn10 are shown in light pink and magenta, respectively. **A)** In the wtFn10-bound conformation, the backbone of Asp of RGD motif (light pink) made electrostatic contacts with the backbone carbonyl oxygen of Arg216, the sidechain amino group of which forms an H-bond with the carbonyl oxygen of Met165. Further, the backbone -NH of Met165 made H-bonds with the backbone carbonyl oxygens of Val161 and Ser162. **B)** In the hFn10 bound structure, Asp (RGD) (magenta) - Arg216 - Met165 - Val161 interaction distances were shorter, and Met165 did not interact with Ser162. **C)** Similarly, in the wtFn10-bound structure, the sidechain of Asp of the RGD motif (light pink) forms H-bond with the backbone of Asn215, while its sidechain carbonyl oxygen formed an electrostatic contact with the backbone carbonyl oxygen of Tyr164. The backbone nitrogen of Tyr164 forms an electrostatic contact with the backbone of Val161. **D)** In the hFn10 bound structure, the interaction distances of the Asp (RGD)-Asn215-Tyr164-Val161 connection network were significantly reduced to form stronger H-bonds.

The residue interactions at the RGD-binding site (site I) and those connecting site II with site I through SDL seem consistently stronger in the hFn10-bound conformation (inactive) than in the wFn10-bound conformation (active). While the Ser162 sidechain forms an H-bond with Ala263 in the active conformation, such interaction is absent in the inactive structure. The H-bond network among Asn215, Tyr164, and Val161 is stronger in the hFn10-bound conformation, which seems to disrupt the interaction between Met165 and Ser162. As Val161 moves closer to Tyr164, the Ser162 backbone is flipped away from Met165 and Ala263, losing the interactions while engaging with His113 of β-propeller. Both D(RGD)-Asn215-Tyr164-Val161 and D(RGD)-Arg216-Met165-Val161 interaction networks appear to coordinate together and likely regulate the communication between site II and the RGD-binding site. These observations suggest that the interactions of the RGD motif at the primary binding site and network communications between the primary RGD-binding site and site II through the SDL residues are significantly different among the wtFn10- and hFn10-bound structures. A schematic representation of the connection is shown in Fig. 12.

**Fig. 12.**
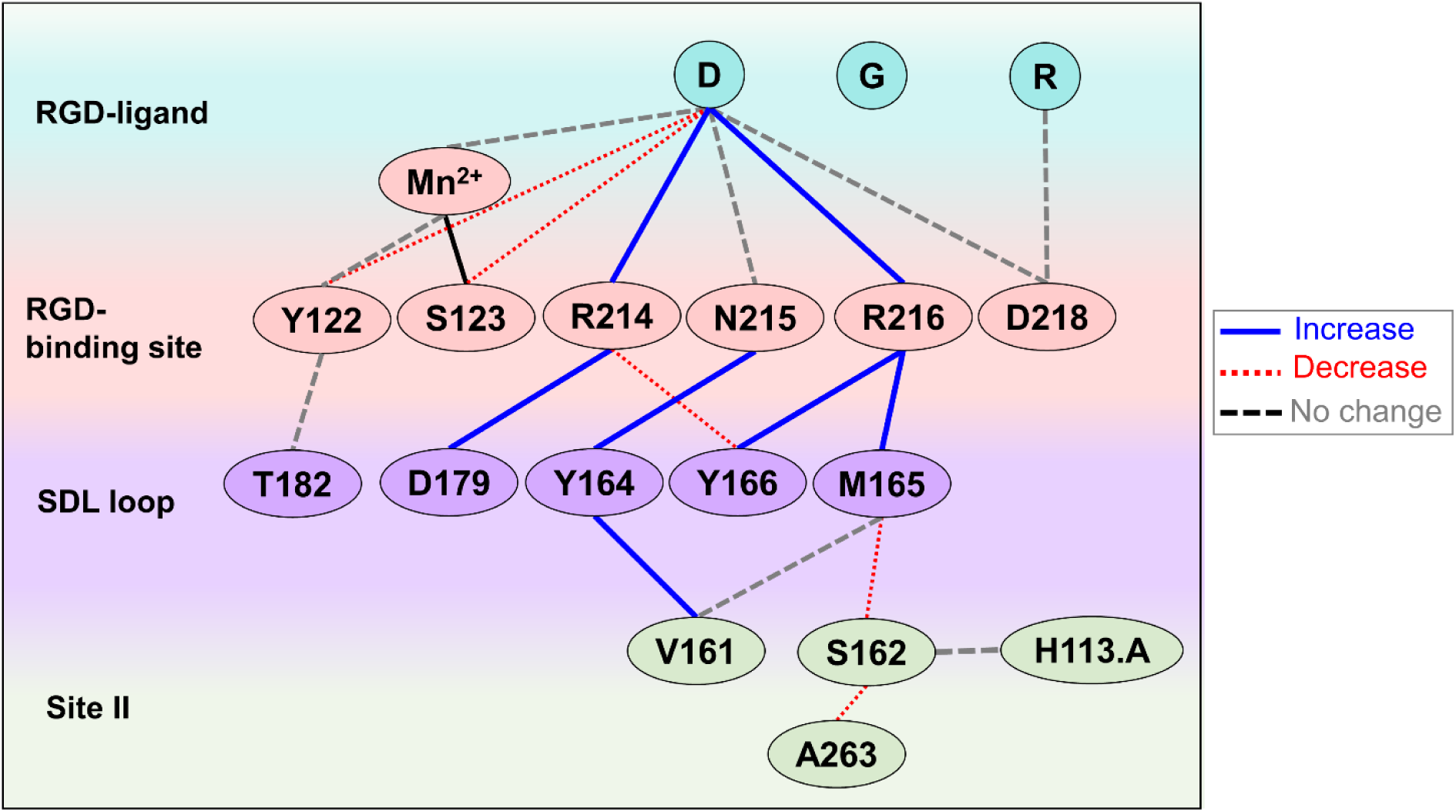
Schematic diagram of the molecular interactions network between the primary RGD-binding site and site II. The network of interactions connecting residues from the RGD-binding site, SDL loop, and site II are shown as lines. The blue and red lines indicate the increase and decrease in the interaction distances, respectively, upon the receptor activation. The black lines indicate the interactions in which the distances remain unaffected in both the active and inactive states of the receptor. Asp of the RGD ligands mainly interacts with three residues Arg214, Asn215, and Arg216 of the primary binding site, which are in turn connected with the site II residues Val161, Ser162, and Ala263 through the SDL loop residues Tyr164, and Met165. SDL loop seems to act as a hub connecting the RGD-binding site and site II.

Comparative analysis of the SDL backbone conformations revealed surprising differences in the torsion angles between the agonist- and antagonist-bound conformations of integrin αVβ3. In the antagonist-bound structures, the backbone peptide bonds between the residue pair Ser162-Pro163 and Ser186-Pro169 are in *cis* conformation while they assume *trans* conformation in the agonist-bound structures (**Fig. 13**). The subtle differences in the H-bond networks D(RGD)-Asn215-Tyr164-Val161 and D(RGD)-Arg216-Met165-Val161 between site II and the primary RGD binding site seem to regulate *cis*-*trans* conformational transition of the Ser162-Pro163 peptide bond. Specifically, Ser162, which is located at site II, has been reported as a functionally important residue for site-II-mediated regulation, and Ser168, located at the SDL loop, engages in direct contact with the β-propeller in the cis conformation. Since *trans*-to-*cis* peptide bond conversion is observed in all the antagonist-bound conformations of integrin (hFn10, TDI-4161, and var-hFn10), the specific residue interactions of both Ser162 and Ser168 might be critical for the proper inhibition of the integrin αVβ3 receptor.

**Figure 13.**
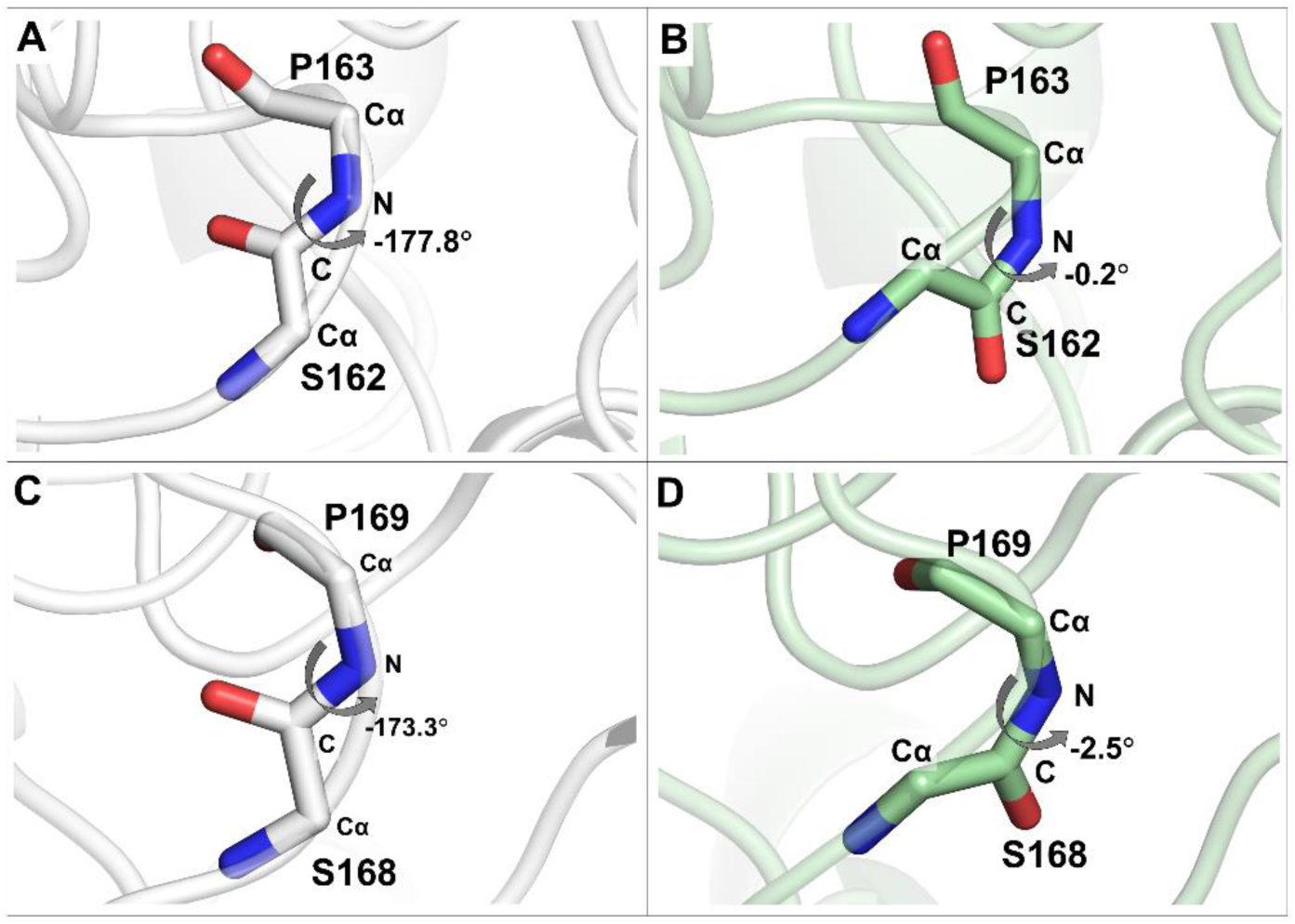
Cis-trans peptide backbone conformational transformation in SDL. The binding of wtFn10 or hFn10 to integrin αVβ3 resulted in differential interactions within SDL and adjacent residues. These differential interactions resulted in specific backbone conformations in the residue pairs Ser162-Pro163. **A-B)** and Ser168-Pro169 **C-D)**. The backbone conformation changes from trans type in the active structure (wtFnFn10, white) to *cis* type in the inactive structure (hFN10, pale green). The C_α_ -C- N- C_α_ torsional angles of the residue pairs, Ser162 -Pro163, in the wtFn10-**A)** and hFn10-bound **B)** structures are −177.8° and −0.2°, respectively. Similarly, the C_α_ -C- N- C_α_ torsional angles of the residue pairs, Ser168 -Pro169, in the wtFn10-**C)** and hFn10-bound **D)** structures are −173.3° and −2.5°, respectively.

### Dynamic cross-correlation network analysis

In addition, to residue-contact analyses using the static crystal structures, the dynamic residue interaction networks were analyzed using graph-based representation in which the residues were defined as nodes while the interactions between the nodes were represented as edges. The network analysis of trajectories obtained from 100 ns MD simulations of the analyzed structures provides semiquantitative measures of the strength of the interactions and dynamic cross-correlations among the interacting residues as communities and critical nodes. It should be noted that the simulation time was chosen short intentionally to minimize drastic changes from the experimental structures. The entire interaction network was divided into smaller communities with densely connected residues as well as critical nodes connecting communities by residues with high betweenness. In general, betweenness is a measure of the degree of communication between specific nodes connecting communities. Therefore, the critical nodes with a higher betweenness may function as regulatory switch points that determine the strength of the interactions.

Firstly, the number of communities formed within the unliganded integrin headpiece (βI and β-propeller domains) was analyzed. Subsequently, a comparative analysis of the liganded (wtFn10- and hFn10-bound) structures revealed that the number of communities in the headpiece decreased upon the binding of ligands. Specifically, the antagonist-bound structures had a less number of communities. This observation suggests that the integrin headpiece movements were highly correlated and densely connected in the presence of antagonists. Further analysis revealed that the decrease in the number of communities was attributed to the decrease of communities primarily in the βI domain and not the β-propeller domain (**Supplementary Table 3**). For instance, the number of communities in the βI domain decreased from 6 in the wtFn10-bound structure to 3 in the hFn10-bound structure. In all the liganded structures, the SDL residues were observed to act as a separate community. However, interestingly, the SDL residues and the site II residues (Ala263) were part of the same community in the unliganded conformations (**Fig. 14A**). In the case of antagonist-bound structures, except for var-hFN10, the SDL residues were separated from site II to form an individual community. This observation suggests that the binding of ligands recruited the SDL residues and strengthened the interactions among them. Among all the structures, the small molecule inhibitor TDI-4161-bound structure had a smaller number of communities (**Fig. 14A**). Although the SDL residues exist as an individual community upon complex formation with hFn10, the remaining βI residues are grouped into two distinct larger communities. This reduced number of communities observed in the βI domain in the presence of antagonists further supports the αVβ3 integrin’s antagonist-dependent “frozen headpiece inhibition” hypothesis [27].

**Figure 14.**
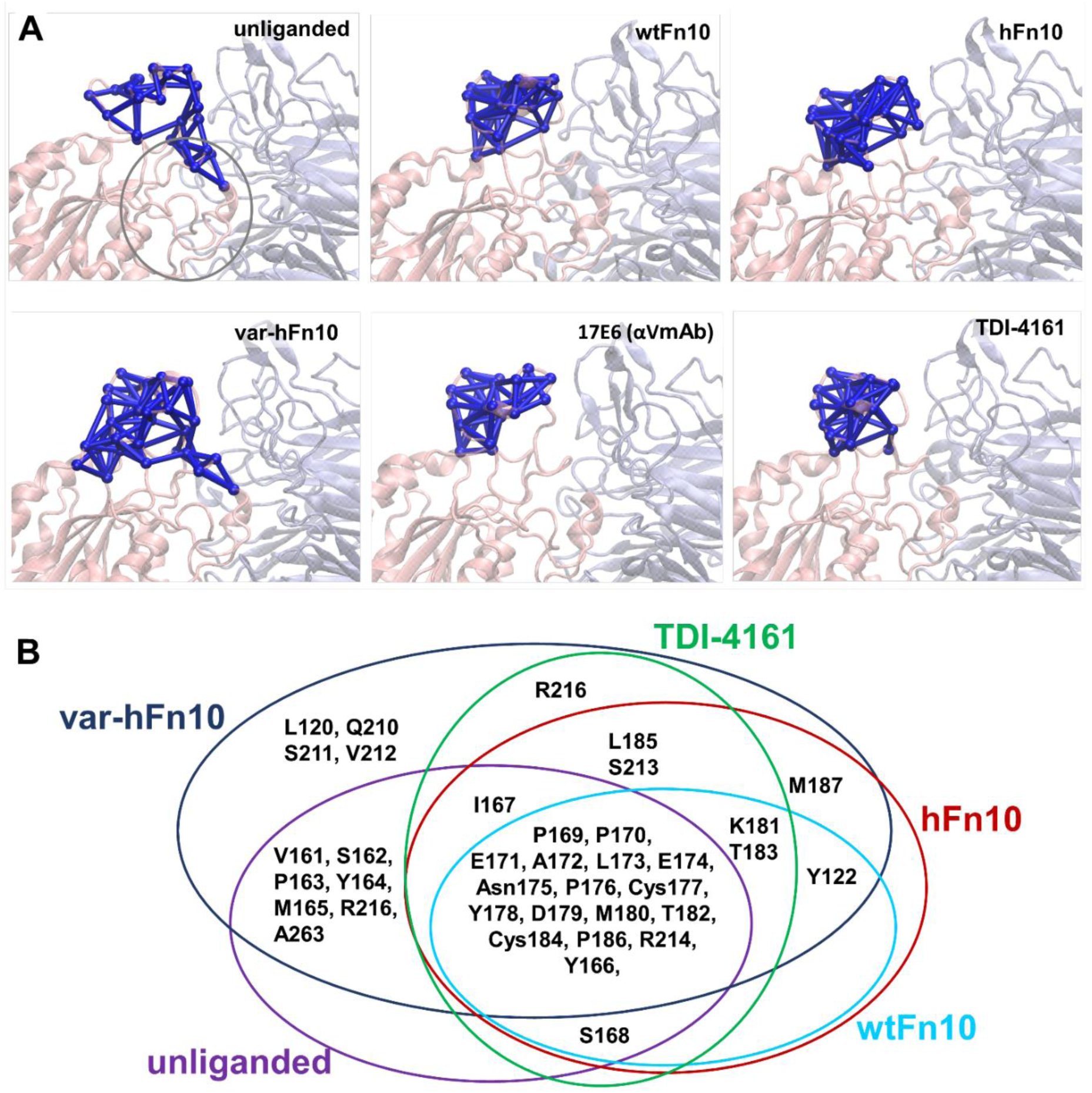
SDL loop residues as communities. αVβ3 headpiece is represented in cartoon mode; the β-propeller domain and βI domain are shown in slate blue and light pink, respectively. Within the network, residues were represented as nodes, and the interactions were represented as edges. The residue community formed by SDL residues is shown in blue. **A)** The community formed by the SDL residues includes site II residues in the unliganded structure. Similarly, in the var-hFn10 bound structure, the community includes the site II residues. In wtFn10-, hFn10-, TDI-4161-, and αV-specific antibody-bound structures, the community is constrained within the SDL residues. **B)** The residue composition of the SDL community for unliganded and liganded structures is primarily formed by SDL residues in unliganded and apo/liganded (wtFn10, hFn10, var-hFn10, and TDI-4161) structures. Circles represent the community and encompass the residues in the community. Circles were colored and labeled differently for each ligand.

Further, analysis of the residue composition of the community formed by the SDL residues shows that the residues Pro169, Pro170, Glu171, Ala172, Leu173, Glu174, Asn175, Pro176, Cys177, Tyr178, Asp179, Met180, Thr182, Cys184, Pro186, Arg214, and Tyr166 were part of a single community and was present in all the structures irrespective of the unliganded- or the liganded-state of the headpiece. In the wtFn10-bound structure, Lys181, Thr183, and Tyr122 were merged with the other residues to form the community. Whereas in the case of hFn10, the community includes seven residues, including Lys181, Thr183, Tyr122, Leu185, Ser213, Met187, and Ile167. In the var-hFn10-bound structure, several other residues were part of the community formed in the hFn10 structure. (**Fig. 14B**). One of the prominent facts observed in the ligand-bound structures was that Tyr122 of the α1 helix becomes a part of the SDL residues community in wtFn10-, hFn10-, var-hFn10-, and αV-specific antibody-bound structures but not in the TDI-4161-bound structure. The placement of TDI-4161 between the SDL and Tyr122 separates the coordinated movement of Tyr122 with SDL, unlike other macromolecular ligands (**Fig. 14B**).

Further analysis of the edges formed at the interface between SDL and the β-propeller domain revealed that in the unliganded conformation, only three SDL residues were found to be in contact with the β-propeller (**Fig.15A**). However, in wtFn10-bound conformation, seven SDL residues Ser162, Pro163, Tyr164, Ile167, Ser168, Pro169, and Asp179 made contact with the β-propeller domain (**Fig. 15B**). In the case of hFn10-bound conformation, only three SDL residues namely, Pro163, Ser168, and Pro169 formed interface edges with the β-propeller, especially Pro163 interacted with several residues of the β-propeller (**Fig. 15C**). In the var-hFn10-, αV-specific antibody- and TDI-4161-bound structures, six (Ser162, Pro163, Tyr166, Ile167, Ser168, Pro169), seven (Ser162, Pro163, Tyr164, Tyr166, Ile167, Ser168, and Pro170) and five (Pro163, Tyr166, Ile167, Ser168, and Pro169) residues were observed to be in contact with the β-propeller, respectively (**Fig. 15C, E and F**).

**Figure 15.**
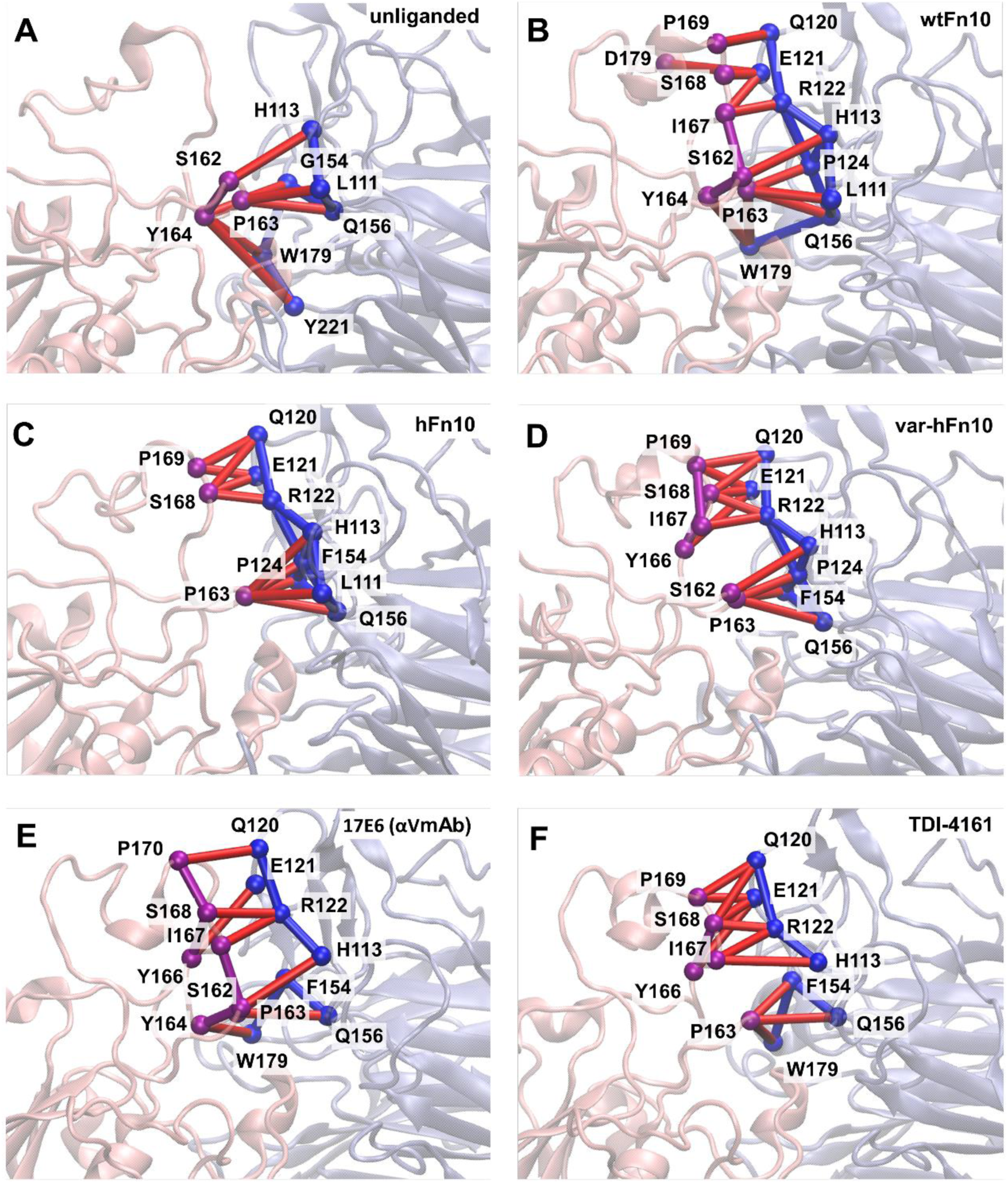
Network connections of SDL residues with β-propeller. αVβ3 headpiece is represented in cartoon mode. The β-propeller domain is shown in slate blue color and the βI domain in light pink. In the network, residues were represented as nodes, and the interactions were represented as red edges. Nodes of the β-propeller and βI domain were shown in blue and magenta spheres, respectively. Edges of the SDL and β-propeller domain were shown for **A)** unliganded **B)** wtFn10, **C)** hFn10, **D)** var-hFn10, **E)** αV-specific antibody, and **F)** TDI-4161 bound structures. **A)** In unliganded structure, three residues of SDL Ser162, Pro163, and Tyr164 form interactions with β-propeller **B)** In wtFn10, seven residues of SDL are in contact with β-propeller. Whereas in the inhibitors such as **C)** hFn10, **D)** var-hFn10, and **F)** TDI-4161 bound structures, a minimum of three residues to the maximum of six residues of SDL interacts with the β-propeller domain.

Also, analysis of the critical nodes at the interface between the βI and β-propeller domains revealed several interdomain edges in the wtFn10-bound structure (**Fig. 15**). In the case of hFn10-bound conformation, only three SDL residues, namely Pro163, Ser168, and Pro169 formed interface edges with β-propeller with an increase in the betweenness of Ser168, which seems critical for maintaining interface interactions. Ser168 appears to act as a bridging residue with a high betweenness in the hFn10-bound conformation. Interestingly, there are no critical nodes between SDL and the β–propeller domain in the wtFn10-bound structure. However, Pro163 and Tyr164 of SDL formed edges with Leu111 and Trp179 of β-propeller. On the contrary, in both hFn10- and αV-specific antibody-bound structures, Ser168 formed critical edges with Arg122 of β-propeller. Both Pro163 and Tyr166 formed critical edges with Gln156 and Asp121 of β-propeller in TDI-4161-bound structure (**Supplementary** Figure 4). A minimum of one critical connection between SDL and β– propeller interface has been observed as an indispensable feature in the antagonist-bound structures.

### Summary and Future Perspectives

Despite enormous efforts by academia and industry, achieving safe inhibition of integrin αVβ3 continues to be an extremely challenging endeavor, with no single FDA-approved drug for clinical use currently. A major stumbling block is the partial agonism produced by current antagonists, resulting in fatal side effects. The mechanistic details of structural and molecular mechanisms contributing to the partial agonistic activity remain poorly understood. However, the availability of vast structural data for integrin αVβ3 bound to various ECM proteins, peptides, small molecule antagonists, and antibodies provides unprecedented details for in-depth analysis to achieve pure antagonism.

In recent years, X-ray crystal structures of integrin αVβ3 bound to the tenth domain of wild-type fibronectin (wtFn10) and its high-affinity mutant version, hFn10, revealed a critical interaction that could entirely arrest the headpiece in an inactive state. A single mutation of S1496W introduced in hFn10 revealed that the pi-pi interaction of the tryptophan sidechain with Y122 of the βI domain could prevent the movement of α1 helix and freeze the receptor in the inactive state. The design of small molecule inhibitors TDI-3761 and TDI-4161, inspired by the hFn10 antagonism, confirmed the validity of such an inhibitory mechanism. Recently, Sen et al. reported the discovery and characterization of an inhibitor MSR03 that can arrest the closed conformation of integrin αVβ3 with a slightly different mechanism than TDI-4161. The design principle was adapted from the previously developed αIIbβ3 inhibitor RUC-4 (Zalunfiban), which interacts with E220 in the MIDAS site and displaces the Mn^2+^ ion, thereby arresting the conformational changes of the integrin[59–62]. RUC-4 is currently under clinical testing for ST-segment elevation myocardial infarction [61, 62]. To achieve a similar inhibitory mechanism for αVβ3, the authors utilized virtual screening, metadynamics-based rescoring, and electron microscopy to identify and optimize a series of compounds that interact with E220[63]. Unlike other inhibitors, MSR03 does not have the carboxylic group that mimics D (of RGD) to interact directly with Y122 and S123 of the α1 helix. Instead, MSR03 was able to inhibit through a direct or metal ion-mediated interaction with E220, which coordinates with Mn^2+^ at MIDAS. The direct interaction of MSR03 with E220 displaced the MIDAS Mn^2+^ ion and locked the receptor in an inactive state. A similar chemical principle was employed to design inhibitors of integrin αIIbβ3 by Springer and his colleagues. The αIIbβ3 inhibitors contain a polar nitrogen atom stabilizing a water molecule that intervenes between Ser123 of α1 helix and Mn^2+^ at MIDAS[64]. The change in the metal coordination is general to integrins, and the expulsion of the MIDAS water is one of the requisites for the conformational transition of the receptor[65].

So far, four successful approaches for pure antagonism of integrin αVβ3 have been reported: 1) prevention of the physical movement of the α1 helix[27, 29], 2) interaction with E220, the residues coordinating with Mn^2+^ at MIDAS[59, 63], 3) stabilization of the water molecule that intervenes between Ser123 of α1 helix and Mn^2+^ at MIDAS[64] and 4) monoclonal antibody-based approaches[37, 38, 66]. The αVβ3 inhibitor TDI-4161 can arrest αVβ3 in the closed conformation by obstructing the movement of the α1 helix through the π-π stacking interaction with the sidechain of Tyr122[29]. Our analysis shows that the substitution of tryptophan or tryptophan-mimic alone is not sufficient to exhibit pure antagonism. In the case of TDI-4161, the indole ring forms an additional π-sulfur interaction with the sidechain of M180 from the SDL loop, facilitating the stabilization of the bound inhibitor. Antagonist hFn10 and var-hFn10 were also observed to interact with the backbone of M180 of SDL. However, peptides such as cilengitide, knottin 2.5D, and hFn10 failed to produce complete antagonism despite having Trp substitution next to the RGD motif. Notably, these molecules do not form additional interactions stabilizing the tryptophan interaction with Tyr122. Successful physical obstruction inhibits the formation of H-bond interactions between the backbones of Tyr122 and Ser123 and facilitates the placement of the water molecule coordinating with the MIDAS Mn^2+^ ion. The other two approaches are associated with the coordination of Mn^2+^ at MIDAS.

In one approach, the inhibitor directly interacts with E220 and displaces the MIDAS Mn2+ ion. Whereas in the second approach, a polar nitrogen atom of the inhibitor preserves a water molecule coordinating with the MIDAS Mn^2+^ ion[64]. Analysis of the crystal structures of the closure-stabilizing αIIbβ3 inhibitors reveals that antagonists can form an H-bond with the backbone of Tyr122 and not with Ser123, suggesting that the backbone interaction with Tyr122 is crucial for receptor activation. Currently, antibody-based therapy is the most successful mechanism for inhibiting integrins in clinical practice[66]. To exhibit selectivity, integrin monoclonal antibodies recognize and bind to the SDL loop in the integrin headpiece[38, 66]. The monoclonal antibodies appear to inhibit the receptor by displacing the SDL loop and introducing steric hindrance to the endogenous ligands. The small molecule inhibitor TDI-4161 was also shown to interact with the SDL loop residues, establishing SDL as one of the most critical structural elements that must be included to achieve pure antagonism. Very recently, highly selective mini proteins were designed using *de novo* deep learning methods for αVβ6 and αVβ8 inhibition. The inhibitory mini proteins were able to achieve exceptionally high affinities (picomolar) and > 1000-fold selectivity over other RGD binding integrins by interacting through residues of the SDL[67].

Our in-depth analyses of the active and inactive conformations of integrin αVβ3 suggest that the SDL loop at the βI domain has multifaced functions, including its role in the heterodimer pairing, ligand specificity, activation, and inhibition of the receptor[58]. The functional importance of SDL has been reported in multiple integrin heterodimer subtypes. In α4β1 integrin, swapping the SDL residues of β1 (CTSEQNC; residues 187-193) with corresponding β3 residues (CYDMKTTC; residues 177-184) altered the ligand binding specificity of α4β1[68]. In α4β7, a cation–π interaction between F185 (a β7 SDL residue) and Mn^2+^ at the synergistic metal ion binding site (SyMBS) has been shown to play a vital role in regulating the affinity, signaling, and biological functions[69]. Further, the four clinically approved integrin-targeting humanized monoclonal antibodies, efalizumab (αLβ2), vedolizumab (α4β7), natalizumab (α4β7 and α4β1), and abciximab (αIIbβ3) were also reported to bind and wrap around the SDL residues to exhibit their inhibitory activity[4, 5, 70, 71]. In the case of αVβ3, studies on the structure and mechanism of action of LM609, a monoclonal antibody clinically used for treating several cancers, and αVβ3-targeted radioimmunotherapy, revealed that LM609 binds to αVβ3 headpiece at SDL and thereby sterically hinders the binding of RGD ligands [38]. In addition, crystal structures of inactive αVβ3 in complex with hFn10, varhFn10, and inhibitor TDI-4161 reveal that the ligands are in direct contact with the SDL loop [27]. Recently, 25-hydroxycholesterol (25HC) and 24-hydroxycholesterol (24HC), oxygenated metabolites of cholesterol, have been reported as the first-ever lipid ligands of integrins[17, 57]. Molecular modeling studies of the integrin αVβ3-25HC and αVβ3-24HC complexes revealed that the ligands likely regulate the integrin activity by binding at site II. Both 25HC and 24H interact with the SDL residues and induce significant conformational changes in SDL [17]. These evidence further supports the functional importance of SDL residues involved in determining agonist and antagonist-specific interactions to regulate the functional and conformational dynamics of integrins. Our analyses show that SDL adopts different conformations in the completely-inhibited receptor. The inhibited conformation of SDL appears to strengthen the interactions in the heterodimer interface through *cis* peptide formation, thus adding additional stringency at the heterodimer interface of the receptor. Hence, exploiting the interactions with SDL also seems to be beneficial for integrin inhibition. Moreover, the residue interaction network analysis revealed that the primary RGD binding site and the allosteric site are connected through SDL interactions. Overall, the regulation of the integrin receptor through the primary site and site II by allosteric ligands involves a network of interactions in which a slight difference in the interaction distances determines the activation status of the receptor. More importantly, the fate of the receptor is determined by a combination of small variations in the molecular interactions connected as a network.

The present study summarizes the structural features of the integrin headpiece in atomistic details and advances our mechanistic understanding of integrin activation and inhibition mechanisms. The results provide valuable insights into the mechanism of action of αVβ3 antagonists and may facilitate the development of new therapies for pure antagonism of integrin either through direct targeting of the primary site or through allosteric modulation through site II via SDL as a novel design strategy.

## METHODS

### Integrin αVβ3 structures and interaction analysis

The structures of integrin αVβ3 were retrieved from the RCSB Protein Data Bank (**Table 1**)[31]. Since the cryo-EM structures lacked sidechain atoms, only X-ray crystal structures were used for the initial interaction analysis and molecular dynamics simulations. The structures were prepared using MOE v2015[72]. During preparation, hydrogens atoms were added, missing residues and sidechain atoms were filled. The structures were briefly minimized to remove the steric clashes and bad contacts. The structure preparation was performed with the Quickprep module of MOE[72]. Intra- and inter-molecular interactions (all non-bonded interactions) in the structures were calculated using MOE and Discovery studio visualizer[73, 74].

### Molecular dynamics simulation

MD simulations were performed with GROMACS v2020[75]. The CHARMM-GUI solution builder was used to build simulation systems[76, 77]. CHARMM36 force field was used for proteins, and the small molecule parameters were generated using the CGenFF server embedded within CHARMM-GUI[78]. TIP3P water models were used to solvate the protein molecules in a rectangular box with 10 Å water padding from the edges of the solute molecules[79]. The 0.15 M NaCl salt concentration was used to maintain the physiological pH. The prepared complexes are subjected to 5000 steps of energy minimization and equilibration to maintain the 310 K temperature and 1 bar pressure. During equilibration, position restraints were applied to the protein atoms and gradually removed to ensure proper equilibration of the system. Finally, the production run was performed. All the covalent bonds involving hydrogen atoms were constrained with the LINCS algorithm[80]. Nose–Hoover thermostat and Parrinello-Rahman barostat were used to maintain the temperature and pressure at 310 K and 1atm, respectively[81]. Long-range electrostatic interactions were calculated with a cutoff of 12 Å using the Particle Mesh Ewald method[82]. Simulations were performed with two fs timestep for 100 ns, and the trajectory was saved for every 10 ps. To assess the stability of the interactions in the crystal structures, The structures were subjected to MD simulation simulated for only 100 ns time length with backbone restrained using a force constant of 10 kj/mol. Trajectory analysis was performed using VMD and in-house tcl and python scripts[83].

### Residue network analysis

The Network View plugin in VMD and Carma were used to analyze the molecular contacts and dynamic networks[83–85]. Residue interaction network maps were used to determine how dynamic contacts are altered allosterically. To generate the contact maps, protein residues were defined as nodes, and the edges were defined by two neighboring nodes if the two residues were within a 4.5 Å distance. From the contact maps, the dynamic residue network was generated by weighing the edges by the covariance calculated from the MD simulation (detailed protocol described elsewhere[84]). The edge weight is a measure that is inversely proportional to the calculated pairwise correlation between the nodes. The communities in the interaction networks refer to smaller subgroups within the dynamic interaction network that exhibit dense connectivity. The calculation of residue contacts, cross-correlation critical nodes, and communities was performed using the Network View plugin[83, 84].

