## Supplementary Information for "Structural insights into the molecular recognition of integrin αVβ3 by RGD-containing ligands: The role of the specificity-determining loop (SDL)"

**Supplementary Table 1**. Coordination properties of the metal ions ($Mn^{2+}$ and $Ca^{2+}$) in and around the RGD binding site (βI domain) of the experimental integrin αVβ3 structures

| Site | Properties | Apo | Apo | Apo | Apo | Apo | Cyclic peptide | hFn10 peptide | Knottin 2.5D | Knottin 2.5F | wtFn10 | hFn10 | Var-hFn10 | TDI-416 | mAb |
|---|---|---|---|---|---|---|---|---|---|---|---|---|---|---|---|
| | **PDB ID** | **1JV2** | **1U8C** | **1M1X** | **3IJE** | **4G1E** | **1L5G** | **4MMY** | **6MSL** | **6MSU** | **4MMX** | **4MMZ** | **6NAJ** | **6MK0** | **4O02** |
| | **Metal ion** | $Ca^{2+}$ | $Ca^{2+}$ | $Mn^{2+}$ | $Ca^{2+}$ | $Ca^{2+}$ | $Mn^{2+}$ | $Mn^{2+}$ | $Mn^{2+}$ | $Mn^{2+}$ | $Mn^{2+}$ | $Mn^{2+}$ | $Mn^{2+}$ | $Mn^{2+}$ | $Mn^{2+}$ |
| (MIDAS) | Occupancy | | | | | | 1 | 1 | 1 | 1 | 1 | 1 | 1 | 1 | |
| | B-factor | | | | | | 17 (24.1) | 65.1 (94.4) | 56.2 (47.3) | 46.7 (45.5) | 48.3 (59.2) | 73.4 (87.7) | 37.3 (65.4) | 80.7 (38.9) | |
| | Ligands | | | | | | O4 | O5 | O6 | O6 | O6 | O6 | O4 | O6 | |
| | Valence | | | | | | 0.4 | 1.8 | 1.7 | 2.5 | 2.2 | 2.1 | 1.4 | 2 | |
| | nVECSUM | | | | | | 0.61 | 0.33 | 0.23 | 0.15 | 0.16 | 0.22 | 0.45 | 0.011 | |
| | Geometry | | | | | | Tetrahedral | Octahedral | Octahedral | Octahedral | Octahedral | Octahedral | Octahedral | Octahedral | |
| | gRMSD | | | | | | 34.5 | 17.3 | 20.3 | 20.6 | 18.7 | 23.6 | 15.7 | 17.5 | |
| | Vacancy | | | | | | 0 | 16% | 0 | 0 | 0 | 0 | 33% | 0 | |
| | Bidendate | | | | | | 0 | 0 | 0 | 0 | 0 | 0 | 0 | 0 | |
| (ADMIDAS) | Occupancy | 1 | 1 | 1 | 1 | | 1 | 1 | 1 | 1 | 1 | 1 | 1 | 1 | 1 |
| | B-factor | 61.9(80.8) | 61.9 (59.1) | 56.2 (81.6) | 55.7 (52.2) | | 30.8 (35.3) | 115.6 (138.9) | 86.2 (77.1) | 141 (63.2) | 88.6 (99.6) | 99.8 (122.3) | 59 (96.2) | 210.4 (40.8) | 271.5 133.1) |
| | Ligands | O5 | O4 | O2 | O4 | | O2 | O6 | O5 | O4 | O6 | O6 | O5 | O6 | O5 |
| | Valence | 1.3 | 1 | 0.8 | 1.2 | | 0.3 | 2.3 | 1 | 0.6 | 2.2 | 1.9 | 1.5 | 1.2 | 1.6 |
| | nVECSUM | 0.47 | 0.63 | 0.82 | 0.41 | | 0.85 | 0.25 | 0.48 | 0.47 | 0.26 | 0.36 | 0.47 | 0.27 | 0.4 |
| | Geometry | Octahedral | Tetrahedral | Tetrahedral | Octahedral | | Square Planar | Octahedral | Tetrahedral | Square Planar | Octahedral | Octahedral | Octahedral | Octahedral | Octahedral |
| | gRMSD | 12.2 | 36.1 | 6.2 | 14.1 | | 21.6 | 5.5 | 34.7 | 35.6 | 3.9 | 9.8 | 12.3 | 12.8 | 10.7 |
| | Vacancy | 33 | 0 | 50% | 33 | | 50% | 33% | 0 | 0 | 33% | 16% | 33% | 16 | 33 |
| | Bidendate | 1 | 0 | 0 | 0 | | 0 | 2 | 1 | 0 | 2 | 1 | 1 | 1 | 1 |
| (SyMBS/LIMBS) | Occupancy | | | | | 1 | 1 | 1 | 1 | 1 | 1 | 1 | 1 | 1 | |
| | B-factor | | | | | 302.8 (132.6) | 28.3 (25.1) | 99.5 (90.8) | 163.5 (51.1) | 154.8 (42.7) | 56.1 (58.9) | 103.3 (79.6) | 32.6 (59.5) | 46.5 (41.5) | |
| | Ligands | | | | | O6 | O3N1 | O6 | O5N1 | O6 | O6 | O6 | O6 | O6 | |
| | Valence | | | | | 2.1 | 0.5 | 2.1 | 1.5 | 2.1 | 2.1 | 2.2 | 2.1 | 2.2 | |
| | nVECSUM | | | | | 0.21 | 0.39 | 0.12 | 0.33 | 0.12 | 0.17 | 0.35 | 0.093 | 0.078 | |
| | Geometry | | | | | Octahedral | Tetrahedral | Octahedral | Octahedral | Octahedral | Octahedral | Octahedral | Octahedral | Octahedral | |
| | gRMSD | | | | | 17.70 | 31.4 | 14.4 | 24.9 | 13.7 | 19.4 | 14.8 | 14.8 | 11.5 | |
| | Vacancy | | | | | 0 | 0 | 0 | 0 | 0 | 0 | 0 | 0 | 0 | |
| | Bidendate | | | | | 0 | 0 | 0 | 0 | 0 | 0 | 0 | 0 | 0 | |

**Supplementary Table 2**. Coordination angles of $Mn^{2+}$ ions of RGD binding site in the wtFn10 (liganded-active) and hFn10 (liganded-inactive) bound structures. Coordination angles that drastically differ between the structures are highlighted in red, and the new metal ion coordination angles are highlighted in blue. (Angles are given in degrees )

| | hFn10 (4MMZ) | | wtFn10 (4MMX) | |
|---|---|---|---|---|
| | **Atoms** | **Angles** | **Atoms** | **Angles** |
| **MIDAS** | Ser B121(OG)- Mn-Glu B 220 (OE1) | 112.1 | Ser B 121(OG)- Mn-Ser B 123 (OG) | 93.3 |
| | Ser B 121 (OG)- Mn-HOH B 801 (O) | 86.1 | Ser B 121(OG)- Mn-Glu B 220 (OE1) | 106.5 |
| | Glu B 220 (OE1)- Mn-HOH B 801 (O) | 133 | Ser B 123 (OG)- Mn-Glu B 220 (OE1) | 149.1 |
| | Ser B 121 (OG)- Mn -HOH B 802 (O) | 143.5 | Ser B 121 (OG)- Mn-HOH B 801 (O) | 164.6 |
| | Glu B 220 (OE1)- Mn-HOH B 802 (O) | 104.4 | Ser B 123 (OG)- Mn-HOH B 801 (O) | 76.7 |
| | HOH B 801 (O)- Mn-HOH B 802 (O) | 69.6 | Glu B 220 (OE1)- Mn-HOH B 801 (O) | 78.5 |
| | Ser B 121 (OG)- Mn - Asp C1495 (OD1) | 91.5 | Ser B 121 (OG)- Mn -HOH B 802 (O) | 84.9 |
| | Glu B 220 (OE1)- Mn-Asp C1495 (OD1) | 84.8 | Ser B 123 (OG)- Mn-HOH B 802 (O) | 63.2 |
| | HOH B 801 (O)- Mn-Asp C1495 (OD1) | 139.7 | Glu B 220 (OE1)- Mn-HOH B 802 (O) | 94.7 |
| | HOH B 802 (O)- Mn-Asp C1495 (OD1) | 90.4 | HOH B 801 (O)- Mn-HOH B 802 (O) | 80.1 |
| | Ser B 121 (OG)- Mn - HOH C1701 (O) | 81.7 | Ser B 121 (OG)- Mn - Asp C1495 (OD1) | 92.9 |
| | Glu B 220 (OE1)- Mn-HOH C1701 (O) | 150.3 | Ser B 123 (OG)- Mn-Asp C1495 (OD1) | 79.2 |
| | HOH B 801 (O)- Mn-HOH C1701 (O) | 71.9 | Glu B 220 (OE1)- Mn-Asp C1495 (OD1) | 121.9 |
| | HOH B 802 (O)- Mn-HOH C1701 (O) | 65.4 | HOH B 801 (O)- Mn-Asp C1495 (OD1) | 96.7 |
| | Asp C1495 (OD1)- Mn-HOH C1701 (O) | 68 | HOH B 802 (O)- Mn-Asp C1495 (OD1) | 142.1 |
| **ADMIDAS** | SerB123(O)- Mn-Asp B 126 (OD1) | 72.3 | SerB123(O)- Mn-Asp B 126 (OD1) | 63.2 |
| | SerB123(O)- Mn-Asp B 126 (OD2) | 120.4 | SerB123(O)- Mn-Asp B 126 (OD2) | 124.4 |
| | Asp B 126 (OD1) - Mn- Asp B 126 (OD2) | 60.8 | Asp B 126 (OD1) - Mn- Asp B 126 (OD2) | 61.3 |
| | SerB123(O)- Mn-Asp B 127 (OD1) | 98.4 | SerB123(O)- Mn-Asp B 127 (OD1) | 88.7 |
| | Asp B 126 (OD1) - Mn- Asp B 127 (OD1) | 61.8 | Asp B 126 (OD1) - Mn- Asp B 127 (OD1) | 89 |
| | Asp B 126 (OD2) - Mn- Asp B 127 (OD1) | 90.2 | Asp B 126 (OD2) - Mn- Asp B 127 (OD1) | 87.1 |
| | SerB123(O)- Mn-Met B 335 (O) | 174.4 | SerB123(O)- Mn-Asp B 251 (OD1) | 69.6 |
| | Asp B 126 (OD1) - Mn-Met B 335 (O) | 105.5 | Asp B 126 (OD1) - Mn-Asp B 251 (OD1) | 128.2 |
| | Asp B 126 (OD2) - Mn- Met B 335 (O) | 61.3 | Asp B 126 (OD2) - Mn- Asp B 251 (OD1) | 158.6 |
| | Asp B 127 (OD1) - Mn- Met B 335 (O) | 76.1 | Asp B 127 (OD1) - Mn- Asp B 251 (OD1) | 110.7 |
| | SerB123(O)- Mn-HOH C1702 (O) | 94.6 | SerB123(O)- Mn-Asp B 251 (OD2) | 103 |
| | Asp B 126 (OD1) - Mn-HOH C1702 (O) | 116.6 | Asp B 126 (OD1) - Mn-Asp B 251 (OD2) | 148.5 |
| | Asp B 126 (OD2) - Mn- HOH C1702 (O) | 77.2 | Asp B 126 (OD2) - Mn- Asp B 251 (OD2) | 122.5 |
| | Asp B 127 (OD1) - Mn- HOH C1702 (O) | 165.3 | Asp B 127 (OD1) - Mn- Asp B 251 (OD2) | 61.4 |
| | Met B 335 (O) - Mn- HOH C1702 (O) | 90.9 | Asp B 251 (OD1) - Mn- Asp B 251 (OD2) | 61.3 |
| **SyMBS / LIMBS** | Asp B 158 (OD2) - Mn- Asn B 215 (OD1) | 94.5 | Asp B 158 (OD2) - Mn- Asn B 215 (OD1) | 87.5 |
| | Asp B 158 (OD2) - Mn- Asp B 217 (OD1) | 101 | Asp B 158 (OD2) - Mn- Asp B 217 (O) | 141.8 |
| | Asn B 215 (OD1) - Mn- Asp B 217 (OD1) | 82.3 | Asn B 215 (OD1) - Mn- Asp B 217 (O) | 88.8 |
| | Asp B 158 (OD2) - Mn- Asp B 217 (O) | 171 | Asp B 158 (OD2) - Mn- Asp B 217 (OD1) | 69.1 |
| | Asn B 215 (OD1) - Mn- Asp B 217 (O) | 85 | Asn B 215 (OD1) - Mn- Asp B 217 (OD1) | 110 |
| | Asn B 217 (OD1) - Mn- Asp B 217 (O) | 70.1 | Asn B 217 (O) - Mn- Asp B 217 (OD1) | 76.6 |
| | Asp B 158 (OD2) - Mn- Pro B 219 (O) | 83.4 | Asp B 158 (OD2) - Mn- Pro B 219 (O) | 83.3 |
| | Asn B 215 (OD1) - Mn- Pro B 219 (O) | 166.3 | Asn B 215 (OD1) - Mn- Pro B 219 (O) | 167.3 |
| | Asn B 217 (OD1) - Mn- Pro B 219 (O) | 111.4 | Asn B 217 (O) - Mn- Pro B 219 (O) | 93 |
| | Asn B 217 (O) - Mn- Pro B 219 (O) | 99.2 | Asn B 217 (OD1) - Mn- Pro B 219 (O) | 58.5 |
| | Asp B 158 (OD2) - Mn- Glu B 220 (OE2) | 89.6 | Asp B 158 (OD2) - Mn- Glu B 220 (OE2) | 124.2 |
| | Asn B 215 (OD1) - Mn-Glu B 220 (OE2) | 66 | Asn B 215 (OD1) - Mn-Glu B 220 (OE2) | 81.8 |
| | Asn B 217 (OD1) - Mn- Glu B 220 (OE2) | 147.4 | Asn B 217 (O) - Mn- Glu B 220 (OE2) | 92.7 |
| | Asn B 217 (O) - Mn- Glu B 220 (OE2) | 98.3 | Asn B 217 (OD1) - Mn- Glu B 220 (OE2) | 163.6 |
| | Pro B 219 (O) - Mn- Glu B 220 (OE2) | 100.4 | Pro B 219 (O) - Mn- Glu B 220 (OE2) | 110.6 |

**Supplementary Table 3.** Number of communities formed within β-propeller and βI domains in the dynamics network cross-correlation analysis.

| PDB ID | Number of Communities | β-propeller | β I |
|---|---|---|---|
| 1JV2 | 12 | 0, 3, 5, 6, 8, 10,11 | 1, 2, 4, 7, 9 |
| 4MMX | 13 | 0, 2, 3, 5, 6, 8, 9, 10 | 1, 4, 7, 11, 12 |
| 4MMZ | 11 | 0, 1, 2, 4, 5, 7, 8, 10 | 3, 6, 9, |
| 4O02 | 10 | 0, 1, 3, 5, 6,8, 9 | 2, 4, 7 |
| 6MK0 | 9 | 0, 2, 4, 5, 7 | 1, 3, 6, 8 |
| 6NAJ | 10 | 0, 2, 3, 5, 6, 7, 8 | 1, 4, 9 |

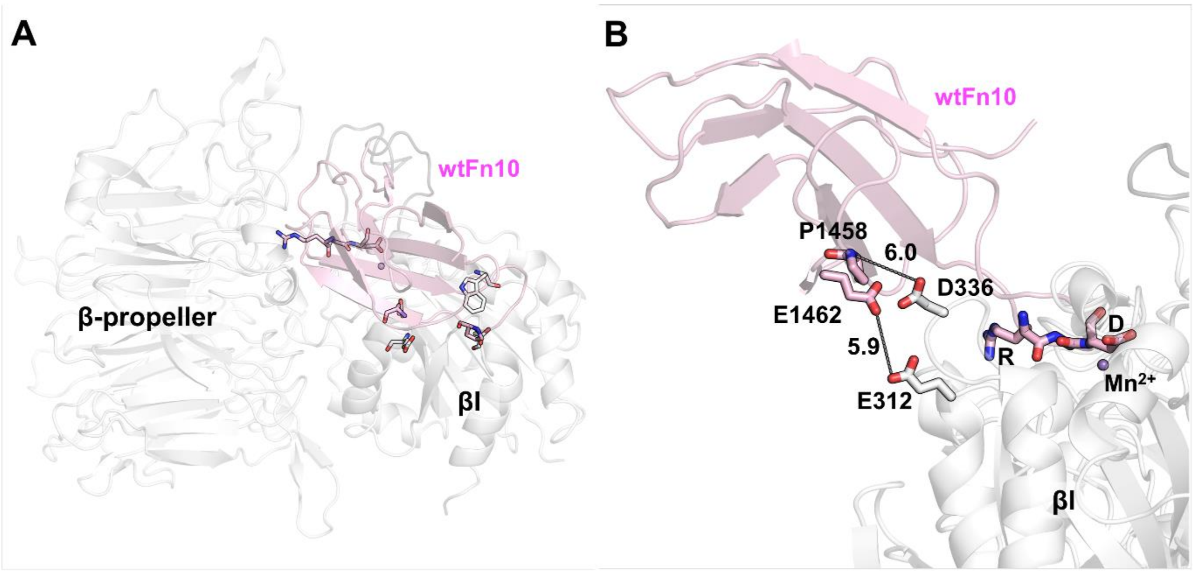

**Supplementary Figure 1. Binding orientation of wtFn10 to integrin αVβ3. A)** The binding orientations of wtFn10 (light pink) is shown in the top view. RGD motif and other residues closer to the α1 and α7 helices are shown as sticks. **B)** Binding interface of wtFn10 (side view) shows that there are no significant interactions with the βI domain of αVβ3 other than those by the RGD motif. The nearest residues (P1458 and E1462) of wtFn10 are ~6 Å away from the βI domain residues (E312 and D336).

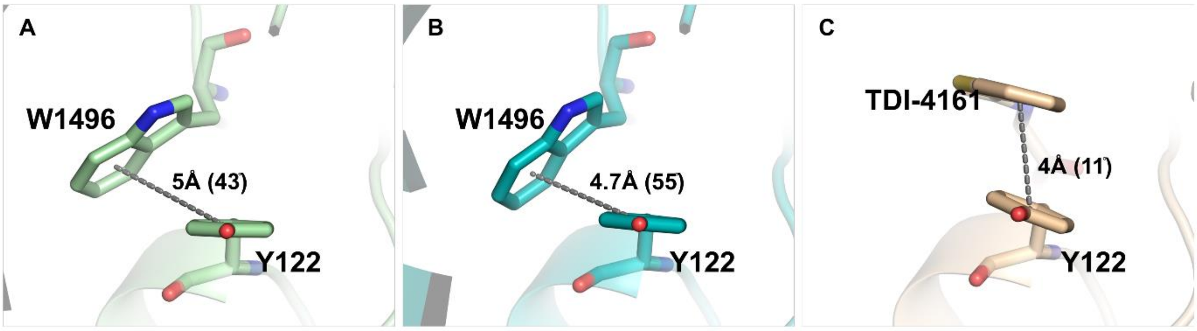

**Supplementary Figure 2. π – π stacking of the Tyr122 of βI domain with integrin αVβ3 agonists.** The π–π interaction is in displaced parallel conformation in **A)** hFn10 and **B)** var-hFn10 bound structures. Trp1496's aromatic ring is situated with regard to Tyr122's plane at angles of 43° and 55° with distances of 5° and 4.7° respectively. **C)** In the case of the TDI-4161 bound structure, the aromatic rings of TDI-4161 and Tyr122 were stacked parallelly at 4Å with 11° angle deviation.

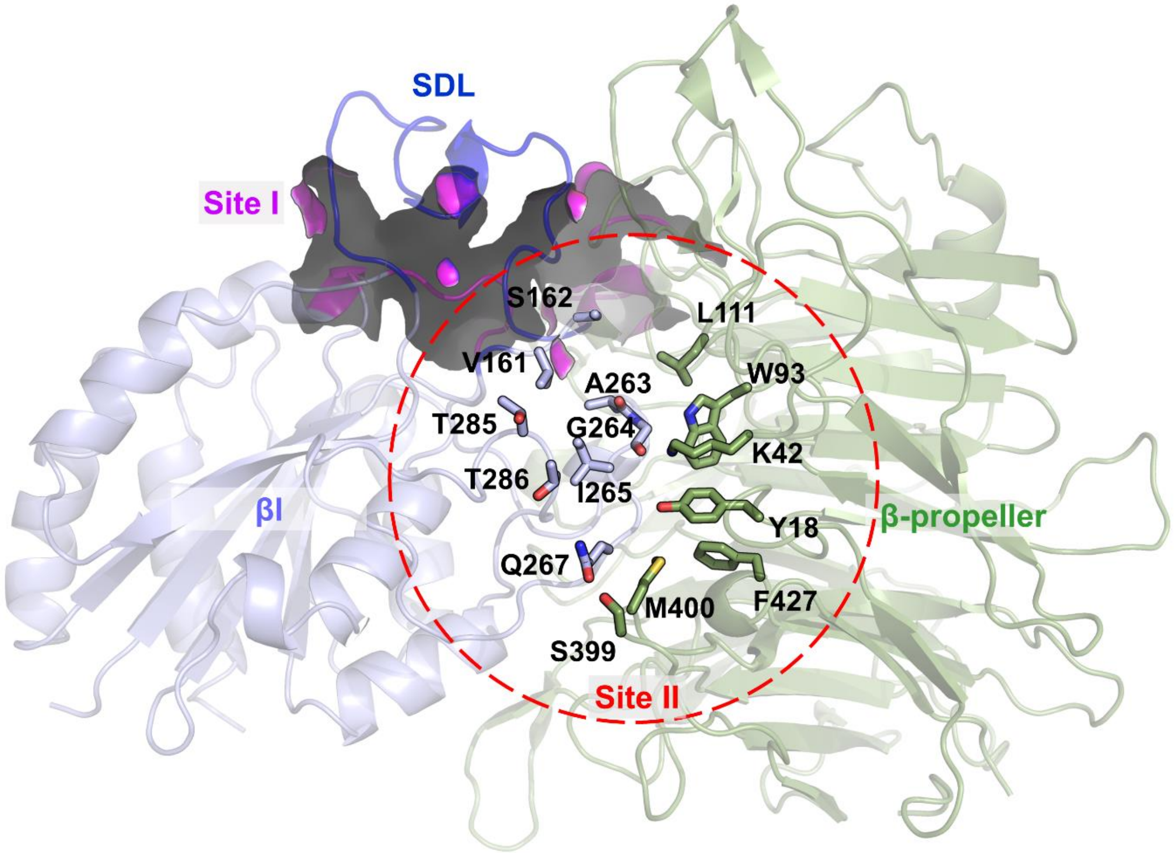

**Supplementary Figure 3. Ligand binding sites at the integrin αVβ3 headpiece.** The critical residues at the allosteric site, Site II (shown as sticks and encircled in red dash lines), is located near the primary RGD binding site (Site I, shown as purple surface) at the αV and β3 interface.

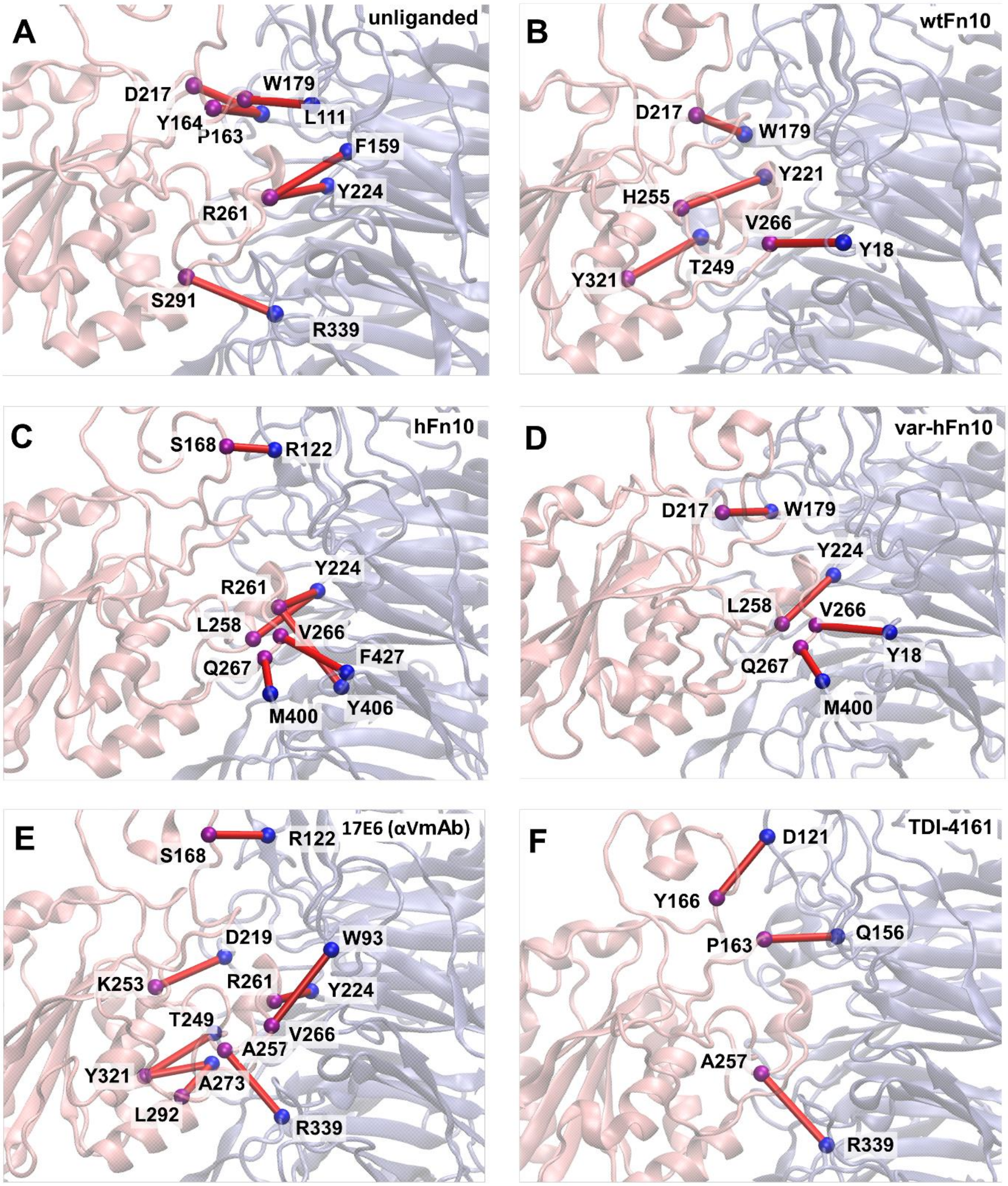

**Supplementary Figure 4. Critical nodes in the αVβ3 headpiece interface region.** αVβ3 headpiece is represented in cartoon mode. The β-propeller domain is shown in slate blue color, and the βI domain is in light pink. In the network, residues were represented as nodes and the interactions were represented as red edges. Nodes of the β-propeller and βI domain were shown in blue and magenta spheres, respectively. Critical nodes in the **A)** unliganded and **B)** wtFn10 bound conformations do not have any critical contact between SDL and the β propeller. Whereas the SDL– β propeller connection formed with **C)** hFn10, **D)** var-hFn10, **E)** αVmAb, and **F)** TDI-4161. Although var-hFn10 binds and has contacts similar to hFn10, var-hFn10 does not have any SDL– β propeller connections in the dynamic environment.
